# Computational Design of Next-Gen Peptide Biopesticides: Targeting the Nicotinic Acetylcholine Receptor in Rice Pests

**DOI:** 10.1101/2025.06.25.661460

**Authors:** Xinyuan Chen, Haojin Zhou, Xueqing He, Jiaqi Wang

## Abstract

The brown planthopper (*Nilaparvata lugens*) threatens global rice production, and conventional pesticides face resistance and environmental challenges. We present a computational framework integrating AlphaFold3-predicted structure of the nicotinic acetylcholine receptor *α*3 subunit (nAChR-*α*3) with multi-force-field molecular dynamics (MD) refinements to design peptide biopesticides. Crucially, we introduce “Dipeptide Probing”, a high-throughput MD-based screening strategy employing 20 phenylalanine-containing dipeptides (F-X) to map dynamic binding sites. Unlike rigid docking or static free energy calculations, this approach captures transient interactions and cooperative binding phenomena, identifying Phe-Met (FM) as the top binder through hydrophobic contacts and multiple hydrogen bonding, while *π*-stacking contributed minimally to complex stability, contradicting conventional paradigms. Additionally, MD simulations revealed an unexpected Aggregation-Induced Hydrophobicity Binding (AIHB) mechanism: FM dipeptides self-assemble via hydrogen bonds, orienting hydrophilic groups toward solvent and exposing hydrophobic surfaces to the target, thereby stabilizing complex formation to the helix bundle surface of nAChR-*α*3. This cooperative behavior, undetectable by docking (e.g., AutoDock Vina failed to predict FM binding) or static energy methods, resolves limitations of reductionist approaches. Our work establishes “Dipeptide Probing” as a generalizable paradigm for dynamic binding-site mapping and underscores AIHB’s potential to revolutionize peptide-based agrochemical design by leveraging emergent intra-/intermulti-molecule interactions.

## Introduction

Rice, rich in carbohydrates,^1^ proteins,^2^ fats,^3^ and a variety of vitamins,^4^ serves as a critical food source for over half the global population.^5^ However, the infestation of rice pests, particularly the brown planthopper (*Nilaparvata lugens*;^6^ BPH), presents a significant threat to the growth and yield of rice,^7^ particularly in China’s Yangtze River basin and southern regions.^8^ BPH sustains and propels its reproductive cycle through the ingestion of rice plant sap. Such feeding behavior not only depletes essential nutrients from rice plants but also facilitates viral transmission (e.g., toothed leaf dwarf disease), leading to stunted growth, excessive tillering, and malformed panicles,^9^ ultimately leading to a significant reduction in rice yield and quality (Figure 1a).

**Figure 1:**
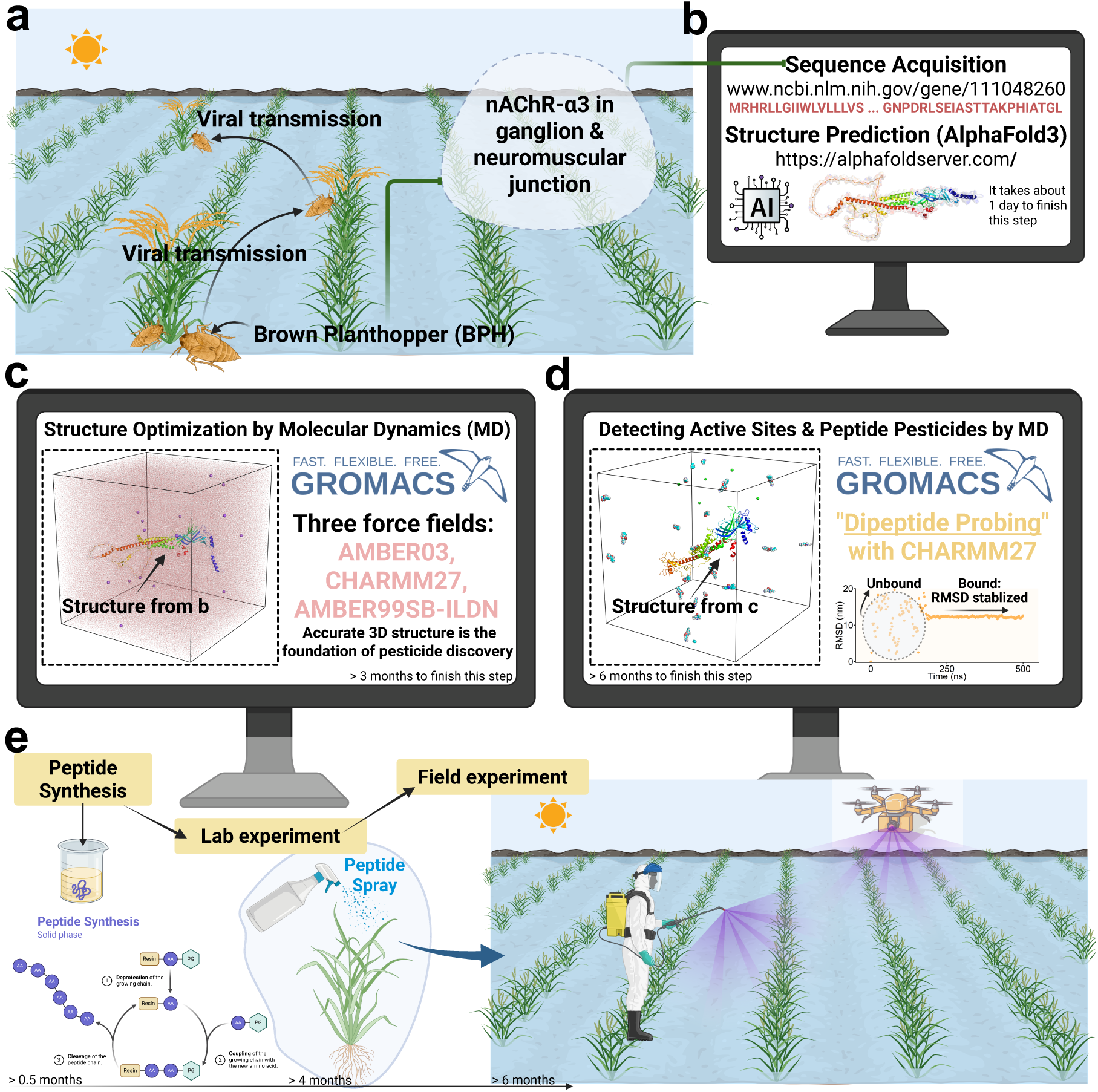
**(a)** The Brown Planthopper (*Nilaparvata lugens*, BPH) is a major rice pest, causing direct feeding damage and potentially transmitting viral pathogens. The nicotinic acetylcholine receptor subunit *α*3 (nAChR-*α*3), predominantly expressed in ganglia and neuromuscular junctions, is a key pesticide target due to its role in synaptic signaling. **(b)** The nAChR-*α*3 amino acid sequence was obtained from a gene database, and its tertiary structure was predicted using AF3. **(c)** The predicted structure was refined via MD simulations in aqueous solution using three force fields (AMBER03, AMBER99SB-ILDN, and CHARMM27). **(d)** The optimized structure was solvated in water solution with 20 distinct dipeptides of the same type for identifying the binding sites (i.e., totally 20 such simulations were performed since 20 types of dipeptides were designed), employing CHARMM27 force field. The stably bound peptides was determined by the RMSD evolution. **(e)** The selected bound peptides will be chemically synthesized and then tested in laboratory experiments to determine how their concentration affects the lethal rate against both nymph and adult BPHs. Subsequently, field experiments will be conducted to validate the efficacy of the peptide pesticides under real-world conditions.

Current strategies for controlling rice pests include agricultural, ^10^ physical,^11^ chemical,^12^ and biological^13^ methods. Chemical control, particularly insecticide application, has been the primary approach for BPH management and remains widely used in modern agriculture.^14^ However, prolonged overuse of chemical insecticides has led to insecticide resistance in BPH populations and accumulation of chemical residues in rice grains, soil and waterways. These issues not only compromise the pest control efficacy but also pose threats to sustainable agriculture and ecosystem health.^15^ Consequently, there is an imperative need to develop eco-friendly alternatives that balance effective pest management with minimal environmental impact.

In light of current agricultural challenges, biopesticides are gaining increasing attention. Among these, peptide pesticides stand out due to their unique advantages, such as high safety profiles,^16^ minimal environmental impact,^17^ high target specificity,^18^ and low toxicity to non-target organisms,^19^ positioning them as sustainable alternatives with significant potential. Peptides, composed of 2-50 amino acids linked by peptide bonds,^20^ play critical physiological roles in living organisms.^21^ Nowadays, peptides have become an indispensable focus of research and development across pharmaceuticals,^22^ agriculture,^23^ cosmetics,^24^ and other fields, with notable success in therapeutics^25,26^ and personal care.^27^ In agriculture, peptide pesticides are emerging as viable replacements for conventional chemical pesticides.^28^ A prominent example is the neuropeptide-based insecticide Spear®.^29^ This innovation was honored with the 2020 U.S. Presidential Green Chemistry Challenge Award for its significant contributions to sustainable agriculture, underscoring the potential and promise of peptide insecticides in modern agricultural practice.

Given the promising potential of peptide pesticides, ongoing research focuses on developing novel agents that target specific proteins in pests. One key target is the *α*3 subunit of the nicotinic acetylcholine receptor (nAChR) in BPH. ^30^ nAChRs are critical neurotransmitter receptors in the insect central nervous system and serve as primary targets for commercial insecticides.^31^ Compounds targeting nAChRs disrupt neural signaling, leading to effective pest control. Developing potential inhibitors targeting the nAChR-*α*3 subunit requires precise structural knowledge of its 3-dimensional (3D) conformation and active sites. In this study, we first predicted the protein structure using AlphaFold3 (AF3) ^32^ (Figure 1b), subsequently we optimized the predicted structure with molecular dynamics (MD) simulations (Figure 1c). However, identifying functional binding sites remained a critical challenge. To address this, we propose a novel “Dipeptide Probing” strategy that systematically maps potential binding regions. This innovative approach involves screening 20 specifically designed dipeptides, each containing phenylalanine (F) paired with one of the 20 standard amino acids (i.e., FA, FC, FD, …, FY). The phenylalanine component serves as a universal *π*-stacking probe, while the variable residues enable comprehensive sampling of diverse binding chemistries. Through extensive MD simulations of each dipeptide-subunit interaction (Figure 1d), we identified key binding residues and potential active sites based on stable interaction patterns. To validate efficacy in real-world contexts, the selected dipeptides will be chemically synthesized and tested in controlled lab studies followed by field experiments (Figure 1e).

In summary, our study achieves three significant advances: (1) High-resolution structural elucidation of the nAChR-*α*3 subunit, resolving its atomic architecture (including critical binding pockets and conformational dynamics) to enable rational design of next-generation peptide pesticides. This structural blueprint directly addresses a key bottleneck in targeted pest control, i.e., the lack of precise molecular targets for biocompatible insecticides; (2) Development of a generalizable “Dipeptide Probing” methodology that universally maps protein active sites. By systematically screening 20 custom dipeptides via MD simulations, we decode binding landscapes with unprecedented efficiency. This paradigm combines peptide versatility with computational scalability, accelerating target discovery for both agricultural and therapeutic applications; (3) Unraveling a new binding mechanism termed Aggregation-Induced Hydrophobicity Binding (AIHB), where peptide aggregation dynamically generates a hydrophobic surface complementary to the target protein. This mechanism, undetectable by static docking or free energy calculations, reveals a fundamentally new paradigm for molecular recognition. Its discovery enables the rational design of peptide therapeutics targeting previously “undruggable” protein surfaces reliant on induced hydrophobicity.

## Methods

### 2.1 Sequence and AF3-Predicted Structure of nAChR-***α***3

The amino acid sequence of the nAChR-*α*3 subunit was retrieved from the UniProt database (www.ncbi.nlm.nih.gov/gene/111048260), which is composed of 579 residues. Its initial structure was predicted using AF3, though we note that AF3 predictions, while providing a valuable starting point, may have limited accuracy. Therefore, we performed MD simulations to further refine the structure (Section 2.2).

### 2.2 Structure Optimization by MD

The accuracy of MD simulations largely hinges on the chosen force field. In this research, we performed all-atom MD simulations with three widely validated force fields, e.g., AMBER03,^33^ AMBER99SB-ILDN,^34^ and CHARMM27,^35^ to optimize the AF3-predicted structure using GROMACS.^36^

The predicted structure was solvated in a cubic box with length of 22 nm, with a minimum distance of 1.0 nm between the protein surface and all box boundaries to prevent artificial periodicity-induced interactions. The system is hydrated using the TIP3P water model^37^ with 350,679 water molecules, resulting in a solvent density approximately 1 g/cm^3^. The system was neutralized with 15 Na^+^ ions to balance the protein’s 15 negative charges.

Energy minimization was performed using the steepest descent algorithm to relax steric clashes and unfavorable contacts. A maximum of 5,000 steps were allowed, with possible termination if the maximum force on each atom fell below 1.0 kJ·mol*^−^*^1^·nm*^−^*^1^. Non-bonded interactions were treated with a cutoff of 1.0 nm for both Coulombic and van der Waals interactions, using a grid-based neighbor search. Long-range electrostatics were approximated with a simple cutoff scheme. Periodic boundary conditions were applied in all directions, and no bond constraints were enforced to allow full flexibility.

Subsequently, the system was equilibrated under NVT ensemble (i.e., constant number of particle, volume, and temperature = 300 K) for 20 ns using a leapfrog stochastic dynamics integrator with a 2-fs timestep. Temperature coupling was applied separately to the protein and solvent groups using the velocity-rescale thermostat with a relaxation time of 1 ps. Finally, a 500-ns production simulation was conducted under NPT ensemble (i.e., constant number of particle, pressure = 1 bar, and temperature = 300 K) using the leapfrog integrator with a 2-fs timestep for a total of 250,000,000 steps (i.e., 500 ns). The velocity-rescale thermostat was also applied for temperature controlling and pressure was regulated at 1 bar using the Parrinello-Rahman barostat with isotropic coupling. In both NVT and NPT simulations, non-bonded interactions were computed using a Verlet cutoff scheme with a 1.1 nm cutoff for both Coulombic and van der Waals interactions. Long-range electrostatics were treated with Particle Mesh Ewald, and the neighbor list was updated every 40 steps via a grid-based search. All bonds involving hydrogen atoms were constrained using the LINCS algorithm to enable the 2-fs timestep. During the NPT simulations, trajectory snapshots were saved every 2 ns, while system energies were written every 2 ps.

Based on the output frames and energy, we employed the potential energy (*E_p_*, with “gmx energy” command), root mean square deviation (RMSD, with “gmx rms” command), radius of gyration (R_g_, with “gmx gyrate” command), room mean square fluctuation (RMSF, with “gmx rmsf” command), and Gibbs free energy landscapes (GFEL, with “gmx sham” command), to characterize the thermodynamic and kinetic evolution of the AF3-predicted structure. The data and figures were processed with Origin and PyMOL.

### 2.3 Detecting Active Residues with “Dipeptide Probing”

It is noted that in the peptide-protein complex, approximately 50% of *π*-stacking interactions involve aromatic rings of phenylalanine in both the protein targets and ligands. ^38^ Leveraging this principle, we systematically designed 20 dipeptides (FA, FC, FD, FE, FF, FG, FH, FI, FK,FL, FM, FN, FP, FQ, FR, FS, FT, FV, FW, FY), each containing F paired with one of the 20 standard amino acids. These dipeptides serve as molecular probes to comprehensively map interaction sites on the nAChR-*α*3 subunit through MD simulations, enabling identification of key binding residues and potential pockets as well as subsequent development of biocompatible peptide-based pesticides targeting BPH.

To characterize the peptide-protein interactions, we conducted MD simulations with optimized protein structure using the CHARMM27 force field. Each simulation box (with edge length of 20 nm and periodic boundary condition) contained one nAChR-*α*3 protein and 20 peptides of the same type, solvated in explicit TIP3P water and neutralized with proper amount of ions. Similar to the structure optimization process, we implemented an energy minimization - NVT - NPT simulation scheme using parameters consistent with structure optimization. A total of 20 independent simulations were conducted, with each simulation running up to 500 ns in NPT ensemble for each one type of dipeptide.

To identify stably bound peptides, we analyzed RMSD of each peptide relative to the protein. Unlike binding free energy calculations, which are computationally expensive and sensitive to force field parameterization, RMSD provides a direct, cost-effective metric for binding stability. A peptide exhibiting equilibrated RMSD indicates stable binding, whereas unbound peptides display continuous fluctuations (Figure 1d). By prioritizing dynamic stability over static energy estimates, RMSD analysis offers a practical, high-throughput alternative to traditional free energy methods for preliminary peptide screening.

## Results and Discussion

### 3.1 Structure Comparison: AF3 vs. MD

The prediction accuracy of AF3-derived nAChR-*α*3 structure was assessed using **p**redicted per-atom **L**ocal-**D**istance **D**ifference **T**est (pLDDT) values, with results visualized in Figure 2a (colored structure) and 2b (score distribution). The model exhibits high-confidence regions (pLDDT ≥ 90, blue; 40.75% of atoms) predominantly in the protein core and stable secondary structures, such as *α*-helices and *β*-sheets. Medium-confidence regions (70 ≤ pLDDT *<* 90, light blue; 32.17%) localize to flexible loops and partial helix segments, suggesting areas potentially involved in dynamic functional processes like ligand binding.

Notably, low-confidence regions (pLDDT *<* 70, yellow-red; 27.08% total, including 18.52% with pLDDT *<* 50) cluster between residues 380 to 520 and in partial helices and random loops, likely representing either functionally important conformational flexibility or prediction uncertainty in these disordered segments. This comprehensive confidence analysis informs subsequent MD refinement by identifying both structurally robust regions and potentially dynamic domains requiring special attention during simulation.

To enhance the accuracy of the predicted 3D structure, particularly in low-confidence regions, we performed MD simulations with three force fields, i.e., AMBER03 (Figure 2c), AMBER99SB-ILDN (Figure 2d), and CHARMM27 (Figure 2e). The simulations revealed significant conformational changes during the initial 20 ns NVT equilibration phase, as evidenced by substantial RMSD values of the 0-ns structure (i.e., optimized structure under NVT ensemble) relative to the AF3 reference structure, specifically, 22.376 Å for AMBER03, 11.716 Å for AMBER99SB-ILDN, and 15.638 Å for CHARMM27 force field. Comparative analysis of structures at 0 ns, 200 ns, and 500 ns under NPT ensemble showed that while all force fields induced major structural rearrangements during equilibration, the systems reached convergence by 200 ns with minimal further changes observed at 500 ns. The AMBER99SB-ILDN exhibited the most pronounced structural deviations, suggesting greater flexibility in accommodating conformational changes. Notably, the simulations demonstrated that low-confidence regions (particularly random loops) were refined into more compact configurations while high-confidence regions maintained their structural integrity, confirming both the dynamic stability of AF3-predicted core regions and the value of MD simulations in improving the accuracy of flexible or disordered regions. More detailed analyses are shown in Section 3.2.

**Figure 2:**
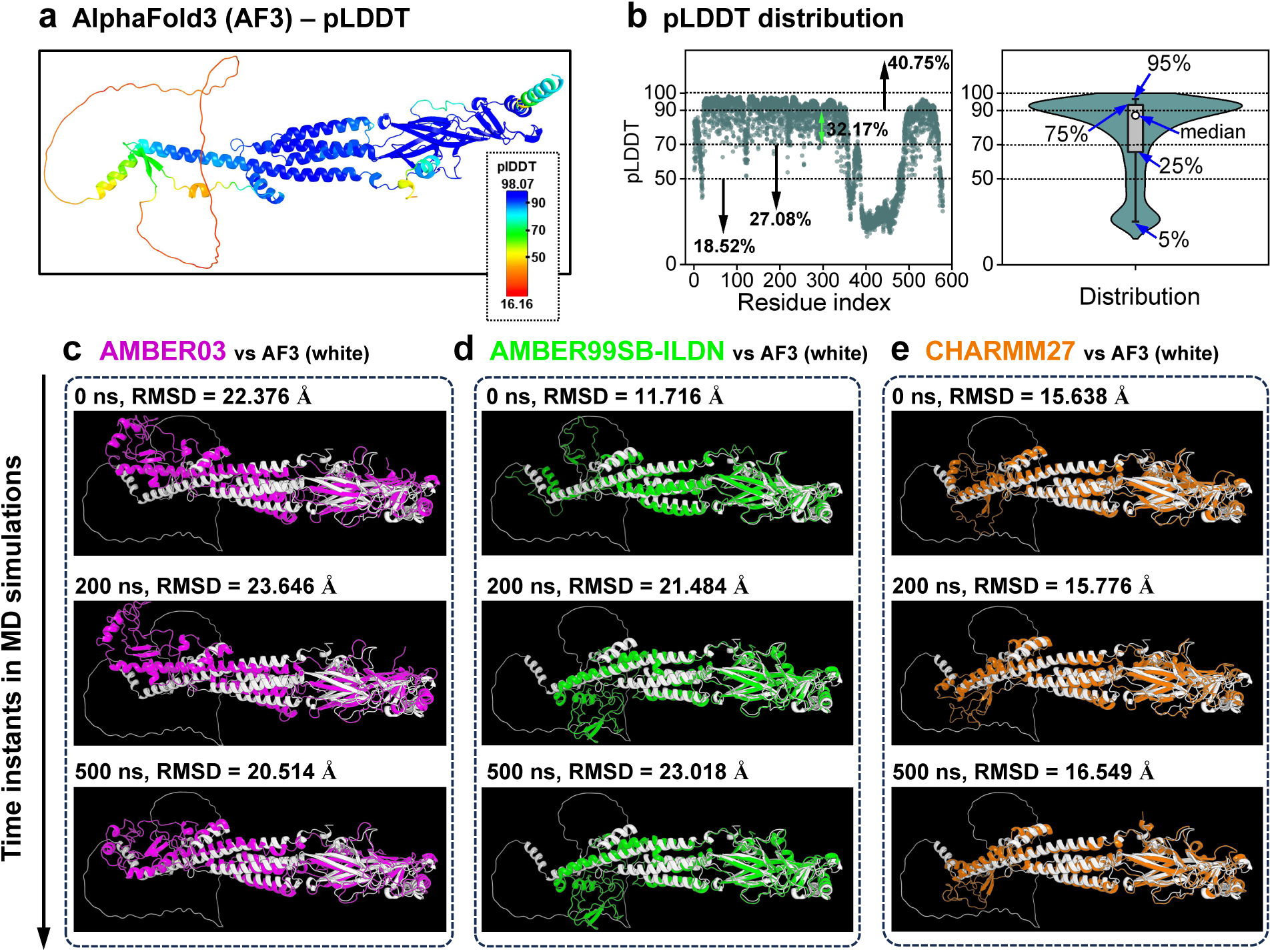
**(a)** AF3-predicted structure of the target protein nAChR-*α*3 subunit, colored by the value of pLDDT confidence scores ranging from 16.16 to 98.07. **(b)** Distribution of pLDDT values across the atoms in the structure (*x* axis indicates which residue the atom belongs to), with 18.52% of atoms exhibiting low confidence (pLDDT *<* 50), 27.08% showing moderate confidence (pLDDT *<* 70), 32.17% falling within high-confidence range (70 ≤ pLDDT *<* 90), and 40.75% displaying very high confidence (pLDDT ≥ 90). A violin plot is included to further illustrate the distribution. **(c–e)** Structural comparisons between the AF3-predicted model (white) and MD simulations during NPT simulations at 0 ns (0 ns-structure indicates one equilibrated by NVT simulation), 200 ns, and 500 ns using the AMBER03 (**c**, purple), AMBER99SB-ILDN (**d**, green), and CHARMM27 (**e**, orange) force fields. The RMSD between the predicted and simulated structures at each time instant was computed using PyMOL to quantify structural differences, with “cutoff” set to 20 Å using 5 outlier rejection cycles. “cutoff” defines the maximum allowed distance in Ångströms between equivalent atoms to be considered a “match” during alignment. Totally 4582 out of 4623 protein backbone atoms were aligned.

### 3.2 Structural Evolution in MD Simulations

MD simulations demonstrated that the *E_p_* profiles for all three force fields decreased significantly within the initial 200 ns, reflecting system relaxation and structural stabilization (Figure 3a). Among the tested force fields, AMBER99SB-ILDN exhibited the most pronounced structural relaxation, with *E_p_* spanning from -17,357.54 kJ/mol to -24,495.55 kJ/mol (Δ*E_p_* ≈ -7,138.01 kJ/mol; 16.0 meV/atom), indicating the most significant structural optimization. While for the other two force fields, the AMBER03 showed an *E_p_* range of -952.18 kJ/mol to -6,621.49 kJ/mol (Δ*E_p_* ≈ -5,669.31 kJ/mol; 12.7 meV/atom), while CHARMM27 displayed intermediate range with *E_p_* values between -13,877.02 kJ/mol and -9,027.96 kJ/mol (Δ*E_p_* ≈ -4,849.06 kJ/mol; 10.9 meV/atom). The enhanced energy decrease observed with AMBER99SB-ILDN may arise from its optimized parameterization, particularly its refined backbone torsional potentials and side-chain rotamer corrections (Ile/Leu/Asp/Asn), which collectively improve conformational sampling while reducing *α*-helical bias present in earlier AMBER versions.

In addition to thermodynamic evolution, we analyzed the kinetic behavior of the system by monitoring the RMSD (Figure 3b), R_g_ (Figure 3c), and RMSF (Figure 3d-f) across all three simulations. The RMSD profiles revealed distinct convergence patterns: while AMBER03 and CHARMM27 stabilized at ∼0.8 nm (*σ* = 0.1 nm, *σ*: standard deviation) after 200 ns, AMBER99SB-ILDN exhibited a more pronounced structural reorganization, with RMSD increasing sharply to more than 2 nm (*σ* = 0.14 nm) during equilibration before plateauing. This substantial deviation, more than double that of the other force fields, reflects extensive conformational rearrangements, particularly in random loop regions with low prediction confidence, as corroborated by that the atoms with pLDDT *<* 50 (Figure 3e) generally exhibit large RMSF values (e.g., atoms within residues numbered approximately 380 to 500). Complementary R_g_ analysis further highlighted these differences: AMBER03 and CHARMM27 maintained relatively constant R_g_ values (∼4 to ∼5 nm), indicating stable global compactness. In contrast, AMBER99SB-ILDN drove a marked compaction, with R_g_ decreasing from ∼4.9 nm to ∼2.5 nm from 200 ns to 500 ns, consistent with its extensive structural adjustments. Collectively, the Δ*E_p_*, RMSD, R_g_, and RMSF metrics demonstrate that MD simulation effectively refine low-confidence regions in predicted structures, and AMBER99SB-ILDN induces the most dramatic conformational changes, likely due to its optimized torsional potentials and explicit all-atom modeling, which enable more significant conformational adjustments toward lower-energy states.

**Figure 3:**
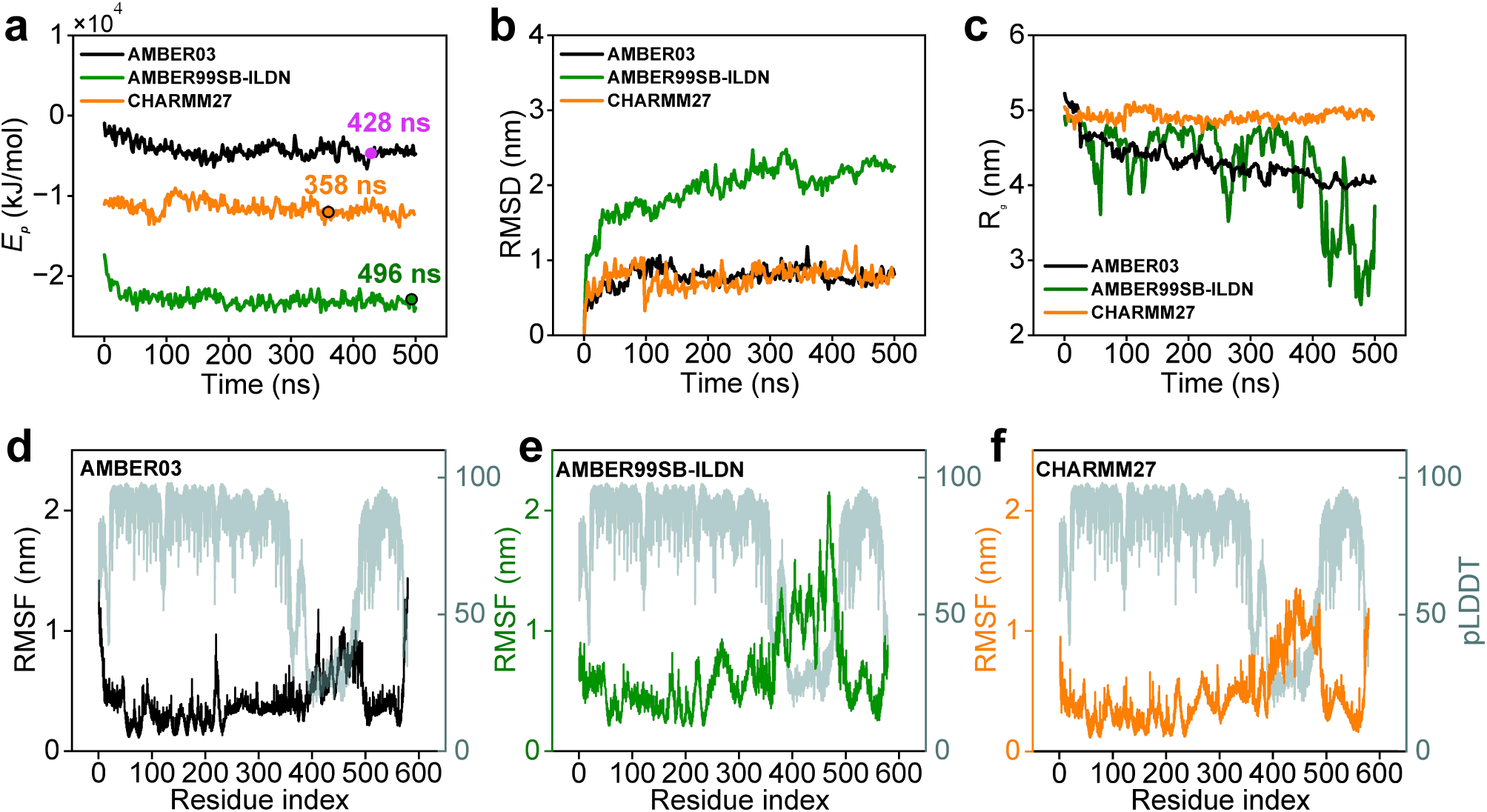
**(a)** Time evolution of potential energy (*E_p_*) during 500-ns MD simulations with NPT ensemble. **(b)** Root mean square deviation (RMSD) of protein atoms relative to the NVT-equilibrated structure, quantifying conformational drift. **(c)** Radius of gyration (R_g_), measuring structural compactness over time. **(d–f)** Root mean square fluctuation (**RMSF, left axis** of each figure) of the lowest-energy conformation with respect to pLDDT (**right axis** of each figure), mapping local flexibility in three force fields to prediction confidence.

### 3.3 Gibbs Free Energy Landscapes of Structures

GFEL analysis of 251 trajectory frames revealed distinct stabilization behaviors across force fields at 300 K (Figure 4): AMBER99SB-ILDN exhibited prolonged energy minimization until 496 ns (RMSD = 22.478 Å), reflecting its capacity for extensive conformational sampling, while CHARMM27 achieved earliest equilibrium at 358 ns (RMSD = 17.097 Å), consistent with its united-atom constraints on flexibility. AMBER03 displayed intermediate behavior (428 ns, RMSD = 19.285 Å), with perturbations primarily in terminal regions. The rapid convergence suggests CHARMM27 effectively maintains structural integrity while permitting limited, localized flexibility, particularly in terminal regions as evidenced by its relatively constant radius of gyration (∼4.8 nm). The force field’s united-atom treatment of aliphatic groups likely contributes to this behavior, balancing computational efficiency with stable sampling of near-native states. CHARMM27’s performance highlights its suitability for systems where preservation of core architecture is prioritized, such as pre-folded proteins or complexes. Therefore, the energy minimized structure of CHARMM27 force field at 358 ns was selected for subsequent “Dipeptide Probing” studies, along with CHARMM27 force field.

**Figure 4:**
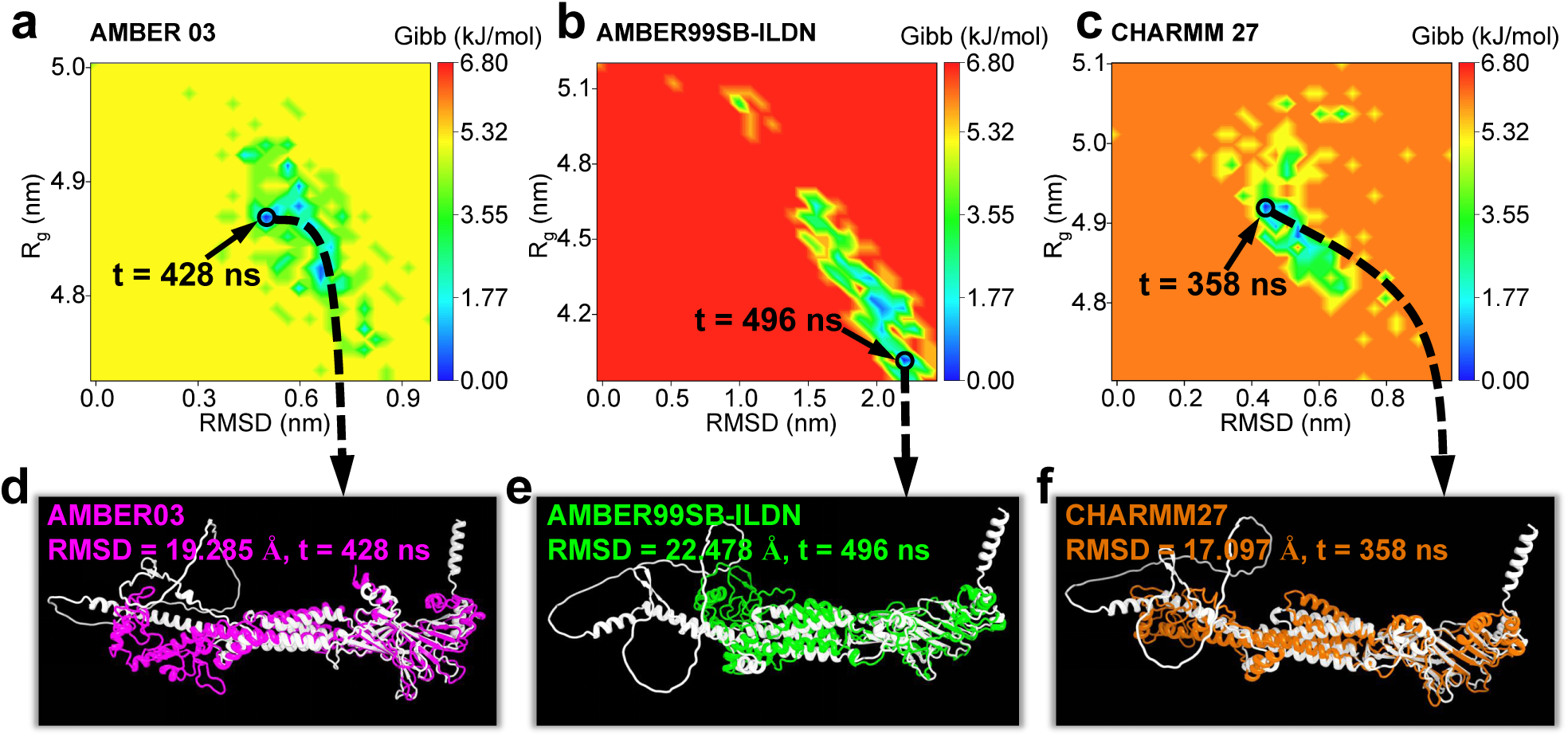
**(a–c)** Gibbs free energy landscapes (GFELs) projected onto principal reaction coordinates (RMSD vs. R_g_), highlighting low-energy conformations found at 428 ns, 496 ns, and 358 ns with AMBER03, AMBER99SB-ILDN, and CHARMM27, respectively. **(d–f)** Structural superposition of the lowest-energy MD-optimized conformation (colored) with the AF3 reference (white) for three force fields. Structural differences are characterized using RMSD output by PyMOL with “cutoff” set to 20 Å, same as in Figure 2c–e.

### 3.4 “Dipeptide Probing” with RMSD

Our “Dipeptide Probing” strategy was deliberately designed to systematically map interaction hot spots of protein. By constructing a comprehensive library of 20 dipeptides - each containing F amino acid paired with one of the standard amino acids, we created a minimal yet maximally informative system to probe all potential interaction types (hydrophobic, aromatic, charged, polar, etc.) at protein interfaces. This approach offers three key advantages over longer peptides: (1) Synthetic accessibility - dipeptides are significantly easier to synthesize and purify at scale compared to tri/tetrapeptides or even longer peptides, facilitating experimental validation; (2) Computational efficiency - the reduced conformational space enables exhaustive sampling of binding modes within achievable simulation timescales; and (3) Interpretability - the binary residue combinations provide unambiguous attribution of specific interactions (e.g., FM’s methionine-driven hydrophobic contacts vs. FR’s arginine-mediated salt bridges). The inclusion of F residue in all dipeptides serves as a conserved aromatic probe (not necessarily a binding “anchor”), while the variable second residue acts as a versatile molecular probe to interrogate complementary binding pockets. This innovative reductionist strategy bridges the gap between single-amino acid mapping and complex peptide discovery, providing atomic-resolution insights into binding pharmacophores while maintaining biological relevance.

To assess peptide-protein binding stability, we employed RMSD of the ligand relative to the protein as a criterion to identify stable dipeptide binders, instead of relying on costly calculations of binding free energies or less accurate docking with predefined pockets/active residues. This approach distinguishes stable binding events from transient interactions by identifying RMSD plateaus, i.e., a stable binding interaction is characterized by the RMSD reaching equilibrium (standard deviation, STD *<* 0.25 nm), in contrast to the fluctuating RMSD patterns typical of Brownian motion. We conducted 20 parallel NPT-ensemble simulations, with each simulation containing 20 peptides of the same type, and we examined 20 RMSD values for each of the 20 dipeptide types, resulting in a total of 400 independent RMSD and STD evaluations (assessed from 300 ns to 500 ns, Figure 5a and 5b, respectively).

**Figure 5:**
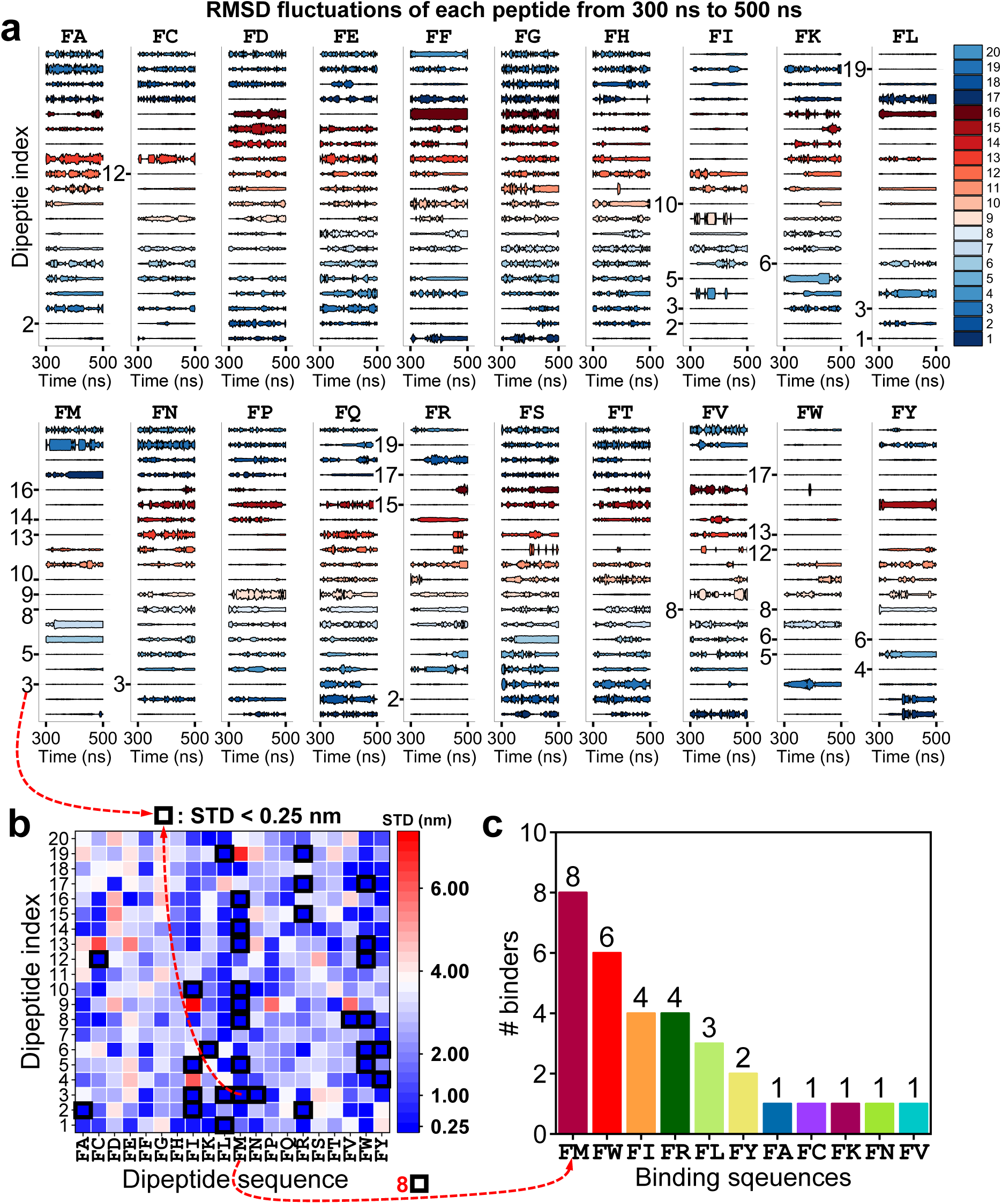
**(a)** Root-mean-square deviation (RMSD) fluctuations of the 20 types of dipeptides from 300 ns to 500 ns. 20 dipeptides of the same type were inserted in one simulation, resulting 20 RMSD fluctuation in each single subfigure. The indices of peptides exhibiting a standard deviation (STD) *<* 0.25 nm over this period are labeled, while others are omitted for clarity. **(b)** Heatmaps of the STD of the 400 dipeptides. Dipeptides with STD *<* 0.25 nm are highlighted with a black border. **(c)** Number of binding dipeptides (out of 20 simulated) per type. Among the 11 binding dipeptides, FM exhibits the highest count (8 binders), while FA, FC, FK, FN, and FV exhibit the lowest count (1 binder each).

Among the 20 types, 11 dipeptide types demonstrated stable binding to the protein of nAChR-*α*3 subunit (Figure 5c). Among these, the FM dipeptide exhibited the highest number of stable bindings, with 8 instances; FW followed with 6 stable bindings. The FI, FR, FL, and FY dipeptides showed 4, 4, 3, and 2 stable bindings, respectively. FA, FC, FK, FN, and FV each had 1 stable binding. In contrast, the remaining dipeptides, including FD, FE, FF, FG, FH, FP, FQ, FS, and FT failed to establish stable interactions with the nAChR-*α*3 subunit. However, it should be noted that if we set the STD criterion larger to 0.3 nm or to 0.5 nm, the type and number of binder may vary and increase. The setting of STD criterion to 0.25 nm in this research ensures robust binder selection.

The exact values of the RMSD of the 400 dipeptides were illustrated in Supplementary Materials. It is evident that prior to binding, the RMSD fluctuates considerably relative to the protein (i.e., pre-binding diffusive). Upon binding, the RMSD rapidly stabilizes to a plateau, with minor fluctuations around the equilibrium position (i.e., post-binding stabilization). This distinct transition in RMSD behavior provides compelling evidence of the dipeptide binding to the protein, thereby validating the stability of the ligand (dipeptide)/target (nAChR-*α*3 protein) complex.

### 3.5 Interaction Analysis between FM and nAChR-***α***3

Since the dipeptide FM was identified as the most potential binder to nAChR-*α*3, we analyzed the interactions of 20 FM dipeptides with nAChR-*α*3 at three time instants (300, 400, and 500 ns) during the MD simulations, as illustrated in Figure 6, with detailed interacting residues shown in Figure 7. Hydrogen bond (HB) interactions were annotated with dashed circles: white circles (labeled A1–A5) indicate stable binding throughout the entire 300 to 500 ns window, while light yellow circles represent less consistent HBs. Among these, tags B, C, and D denote different dynamic behaviors: B-tagged HBs were absent at 300 ns but formed later (400–500 ns), C-tagged HBs disappeared at 500 ns with peptide migration (less stable binding), and D-tagged HBs were less stable and transient, lasting only at 300 ns without persisting beyond 100 ns. These transient HBs may reflect intermediate states during binding or unbinding pathways, suggesting that longer simulations could provide deeper insights into their kinetic roles and the stability of these interactions. Five dipeptides formed stable HBs, with interactions primarily involving charged residues (ARG-481, ARG-482, LYS-342, ASP-518, ASP-330, GLU-205, ASP-63, and ASP-30, Figure 7). The recurrence of certain residues, such as ARG-481 and ASP-518, in HB formation likely stems from their electrostatic complementarity or positioning in key binding pockets, which favors persistent interactions. Notably, peptide A2 exhibited slight instability, shifting rightward over time (indicated by the light blue arrow in Figure 6), leading to changes in interacting residues between 300 and 500 ns. In contrast, peptides A1 and A3–A5 maintained at least one consistent residue in HB formation, suggesting stronger binding. The variability in HB stability underscores the dynamic nature of peptide-receptor interactions, where even minor conformational changes can alter binding modes.

**Figure 6:**
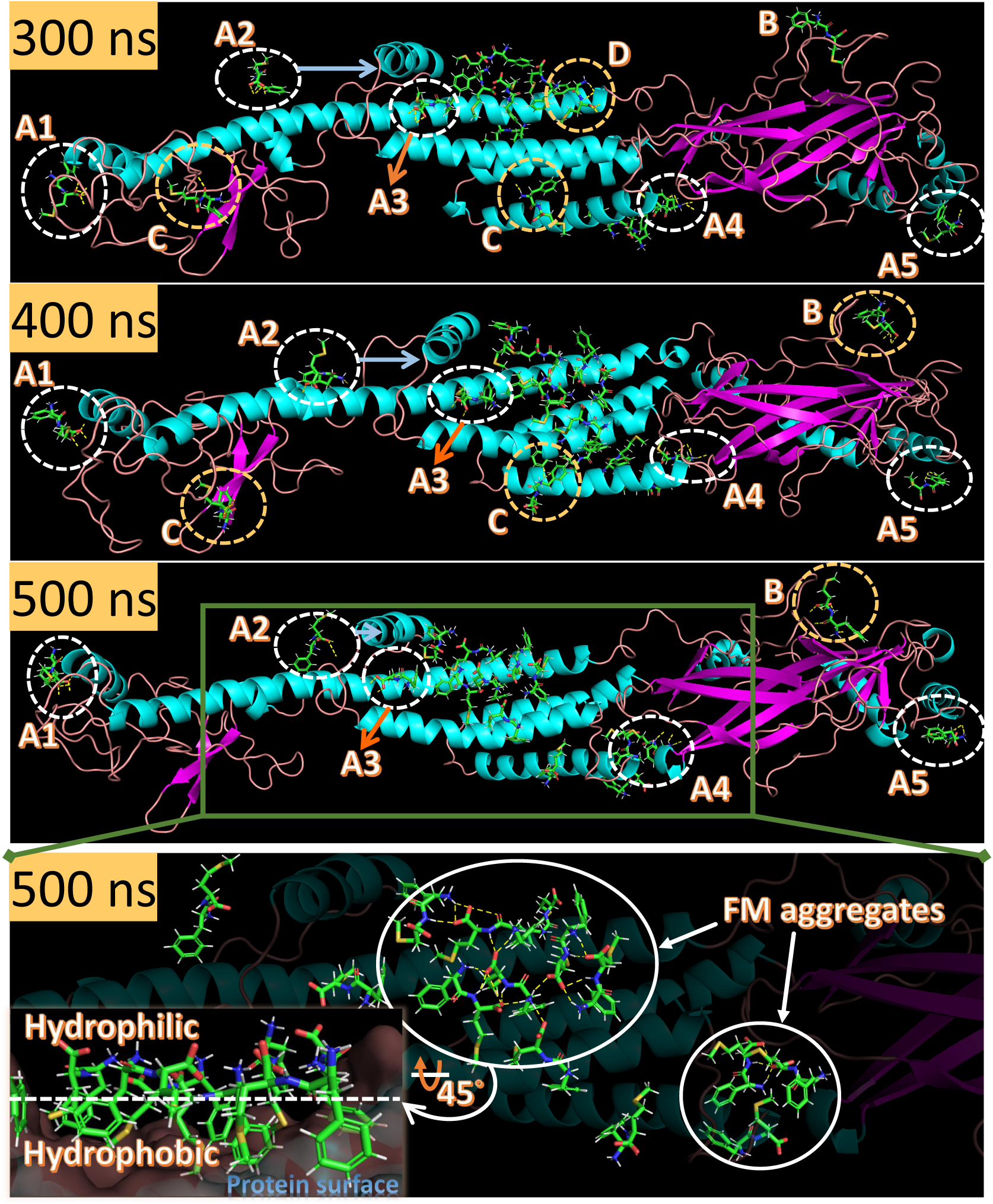
Structural evolution of protein-20 FM complexes from 300 to 500 ns. The peptides which can form hydrogen bonds (HBs) are annotated with dashed circles and are classified into four categories: class A (A1 to A5, rapid stable binding) with persistent HBs throughout the simulation; class B (slow stable binding) forming stable HBs by 400–500 ns after initial absence at 300 ns; class C (unstable binding) with transient HBs (300–400 ns) that dissociate at 500 ns; class D (instant binding) exhibiting brief HB interactions. The magnified 500 ns snapshot (green box) illustrates that the FM aggregation through hydrogen bonding generates a hydrophobic interface that stabilizes peptide binding to protein, with the hydrophilic FM surface orientating toward the water solvent (water omitted for visualization; simulations were fully solvated) and the hydrophobic surface of aggregates faces towards the protein surface. We propose this as an **aggregation-induced hydrophobicity binding (AIHB) mechanism**.

**Figure 7:**
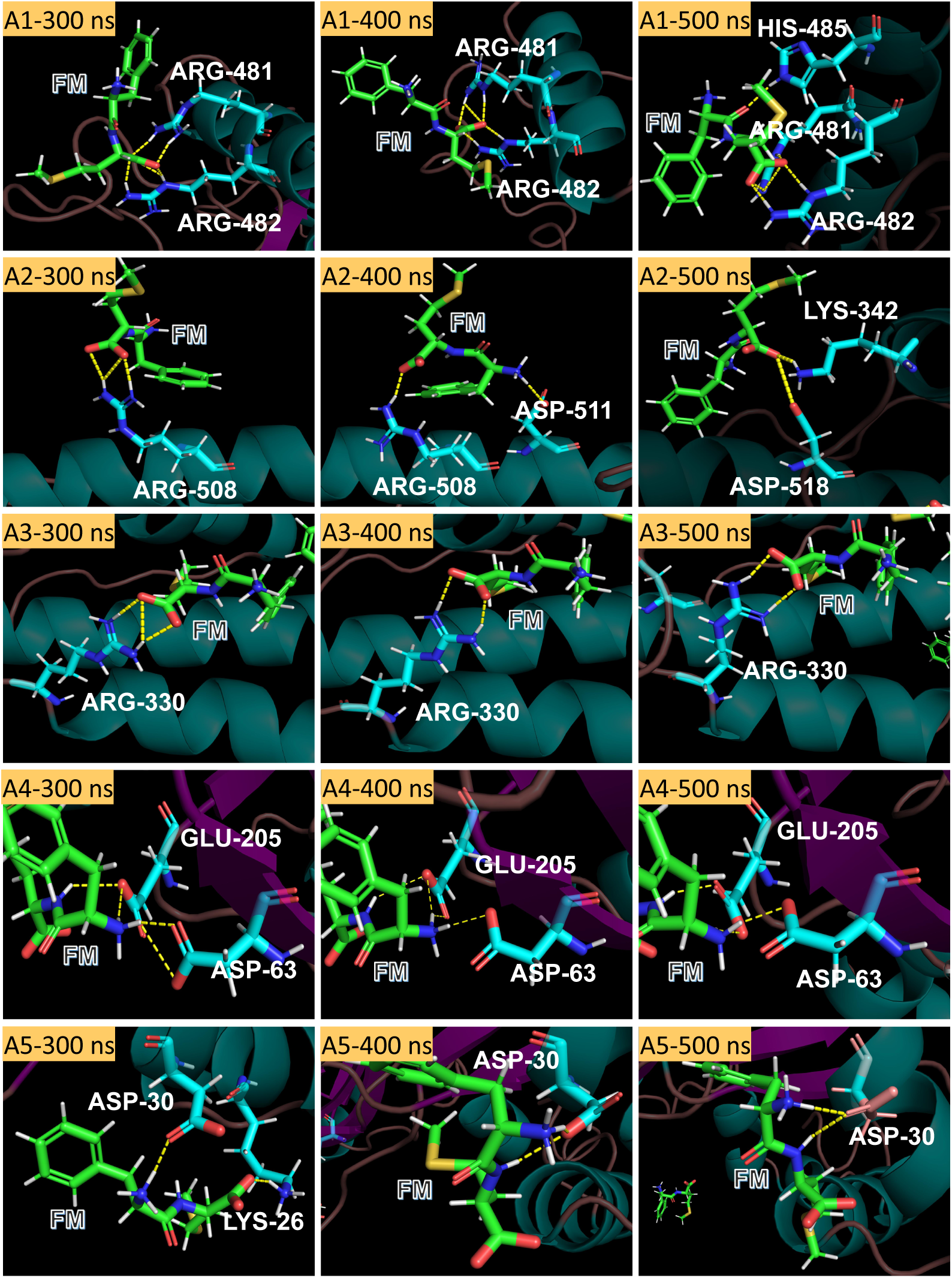
Analysis of interacting residues between the protein and FM dipeptides A1-A5 (as circled in Figure 6) during 300-500 ns molecular dynamics (MD) simulations. MD simulations reveal that all hydrogen bond (HB)-forming residues are charged amino acids and the interactions exhibit a temporal dependent dynamic behavior: Peptide dA1 exhibits lateonset binding with HIS-485 participating only at 500 ns, A2 demonstrates continuously evolving interactions throughout the simulation, and A5 loses LYS-26-mediated HBs after 400 ns. In contrast, A3 and A4 maintain consistent interacting residues across the entire timeframe. These observations collectively demonstrate that dipeptide-protein binding constitutes a complex dynamic process, where transient interaction events, undetectable in static docking analyses, play significant roles in the molecular recognition mechanism.

Interestingly, our preliminary docking predictions using AutoDock Vina failed to identify any binding poses for FM dipeptides, starkly contrasting with the MD simulation results. This discrepancy may indicate the limitations of rigid docking approaches in capturing the dynamic binding behavior observed in MD simulations. Specifically, the flexibility of both the dipeptides and the receptor, along with induced-fit binding mechanisms, likely plays a critical role in stabilizing interactions, factors that are better accounted for in MD than in static docking. These insights underscore the importance of MD in studying peptide-receptor interactions, particularly where docking methods fall short in predicting transient or conformationally dependent binding events.

### 3.6 Aggregation-Induced Hydrophobicity Binding (AIHB)

In addition to traditional single ligand-protein binding interactions, the MD simulations revealed a unique synergistic effect on binding driven by peptide aggregation and hydrophobicity, where FM dipeptides aggregate into stable clusters inducing hydrophobic cluster surface, and binds to the nAChR-*α*3 helix bundle surface. We terms this binding mechanism as Aggregation-Induced Hydrophobicity Binding (AIHB). It should be noted that unlike typical F-contained peptide aggregation driven by *π*-*π* stacking of aromatic groups, these FM clusters form primarily through intermolecular HBs, inducing a remarkably stable orientation: hydrophilic methyl groups face outward to water solvent, while hydrophobic aromatic rings position inward toward the protein interface (Figure 6, last panel). This unique spatial arrangement creates a stable binding interface through synergistic effects of peptide-peptide interactions and optimized hydrophobic complementarity with the target.

The AIHB mechanism represents a significant departure from conventional binding/aggregation paradigms in three key aspects: 1) It demonstrates that stable target engagement can emerge from collective peptide behavior rather than single-molecule interactions; 2) It reveals hydrogen bonding, rather than aromatic interactions, as the driving force for functional peptide aggregation; 3) It establishes a new principle for molecular recognition where self-assembly and binding occur as coupled processes, due to the formed hydrophobic surface of self-assembly. This finding challenges the traditional view of peptides as simple, monomeric binders and suggests that many reported peptide-protein interactions may actually operate through similar cooperative mechanisms that have been overlooked due to limitations/challenges of conventional structural methods. Such behavior is highly relevant for peptide-based drug design, as aggregation-prone sequences may exploit AIHB to achieve prolonged residence times or cooperative binding effects.

### 3.7 Summary of Interactions of Dipeptides and nAChR-***α***3

In summary, the comprehensive analysis of dipeptide binding to the nAChR-*α*3 subunit identified 11 distinct binders, with detailed intermolecular interactions documented in Table 1 and Extended Data Figures 1-16. The binding interface is characterized by charged residues (ARG356, LYS271, ASP527, GLU205, ARG330), aromatic residues (PHE262, TYR263), and polar residues (SER266), which facilitate complex stabilization through extensive hydrogen bonding networks and salt bridge formations. Notably, *π*-*π* stacking measurements revealed intermolecular distances consistently exceeding 5.0 Å (predominantly *>* 6.0 Å), indicating these interactions do not contribute significantly to thermodynamic stabilization of the bound complexes. This observation contrasts with established literature,^39–41^ emphasizing *π*-*π* stacking (typically 3.5 − 4.5 Å) as a primary stabilization mechanism, while our results suggest instead, that hydrophobic nature of aromatic rings may serve as an initial recognition force that orients peptides toward the receptor’s hydrophilic surface prior to formation of higher-affinity HB or electrostatic contacts, but not as a main contributor to the stability of ligand/peptide complex.

**Table 1:** Dipeptides [identified in Figure 5(c)] - residue interaction profiles. Residues with multiple occurrences are formatted in bold.

| Dipeptides | Interacting residues in nAChR-α3 |
| --- | --- |
| FA | **ARG356** |
| FC | **PHE262**, **SER266**, **LYS271**, **ASP527**, ASP63, **GLU205** |
| FI | **ARG330**, LYS140, GLU199 |
| FL | LYS305, GLN548, ARG215, GLU217, **ARG528** |
| FM | **ASP30**, ASP63, **GLU205**, **ARG330**, LYS342, ASP518 |
| FN | **PHE262**, **TYR263**, **SER266**, **LYS271**, **ASP527** |
| FR | HIS328, HIS417, GLU517, LYS520, ASP363, LYS507, GLU510, ASP511, GLU225, **GLU205**, GLU291 |
| FT | ARG365, ASP386, SER398, SER401 |
| FV | **TYR263**, **ARG528**, **GLU205** |
| FW | HIS3, ARG4, **ARG528**, THR535, **ARG356**, **ARG330**, PHE219 |
| FY | **ASP30**, SER34, TYR523 |

Beyond these pairwise interactions, aggregation phenomena through HB were observed for several other dipeptides other than FM (FL, FY, FW; Extended Data Figures 13, 15, 16), wherein hydrophobic moieties associate with the helix bundle surface of nAChR-*α*3 while hydrophilic groups remain solvent-exposed. This spatial organization provides additional validation for the proposed AIHM mechanism, wherein initial aggregate-scale association precedes specific molecular recognition. The AIHM framework resolves an apparent paradox: while individual *π*-*π* interactions lack optimal geometry for stabilization, collective hydrophobic clustering facilitates kinetic trapping of peptides at the receptor interface, creating opportunities for subsequent specific contacts.

For future peptide design, strategic incorporation of bulky hydrophobic residues (e.g., Trp, Phe derivatives) is recommended to amplify both initial binding kinetics (via AIHM) and equilibrium stability (through optimized charge-dipole networks). Such designs should maintain balanced hydrophilicity to prevent nonspecific aggregation while ensuring sufficient surface complementarity with the receptor’s mixed-polarity binding cleft, particularly within the helical bundle region where AIHM-driven organization predominates.

## Discussions and Future Work

Unlike traditional “docking - experiment” validation scheme or static calculation of binding free energy for binding affinity assessment, our study introduces a much more cost-effective “Dipeptide Probing” approach that leverages MD simulations and RMSD stability-based screening to overcome the limitations of conventional docking methods, which fail to capture dynamic phenomena like the AIHB mechanism we proposed. Unlike static docking, our MD simulations revealed how FM dipeptides self-assemble through solvent-mediated interactions and hydrogen bonding to form stable complexes with nAChR-*α*3, demonstrating that peptide binding can emerge from cooperative multi-molecule behavior rather than single-molecule interactions. This finding challenges traditional drug discovery paradigms and suggests that many peptide-protein interactions may involve similar overlooked cooperative mechanisms. The AIHB phenomenon, where peptide aggregates maintain stable binding through optimized hydrophobic complementarity while exposing hydrophilic groups to solvent, opens new possibilities for designing peptide therapeutics but also warrants careful investigation of aggregation propensity to balance efficacy and safety. Future research should explore the generality of AIHB across targets, develop computational or artificial intelligence prediction tools to predict such cooperative binding, and establish design principles to harness this mechanism while avoiding pathological aggregation, ultimately advancing peptide drug/pesticide discovery beyond the constraints of reductionist single-molecule approaches. To comprehensively validate the efficacy and safety of selected dipeptides (e.g., FM, FW) as potential agents for controlling BPH populations, a multi-tiered experimental approach encompassing both laboratory and field studies should be implemented. Initial laboratory bioassays should evaluate the dose-dependent mortality effects on BPH nymphs and adults, while field trials would assess real-world performance across different rice cultivation systems and environmental conditions. Given the ecological importance of pollinators, particularly bees, it is crucial to conduct thorough ecotoxicological assessments to determine any off-target effects on beneficial insects, including contact toxicity tests, feeding deterrence studies, and potential impacts on pollinator behavior and colony health. Furthermore, comprehensive toxicological evaluations must be performed to ensure food safety, including acute and chronic toxicity testing in mammalian models, residue analysis in edible rice grains, and assessment of potential bioaccumulation through the food chain. These studies should be complemented by environmental fate analyses to understand the degradation pathways and persistence of these dipeptides in agricultural ecosystems. The integration of these validation protocols will not only confirm the practical applicability of the dipeptides as eco-friendly BPH control agents but also address potential regulatory requirements for their eventual commercialization and field deployment, while ensuring minimal ecological disruption and maximal food safety standards. We have begun cultivating rice plants (Figure 8) as the foundation for our comprehensive validation studies of the selected dipeptides (e.g., FM, FW), which will systematically progress from controlled laboratory bioassays to field-scale evaluations of BPH control efficacy, environmental safety, and food security implications.

**Figure 8:**
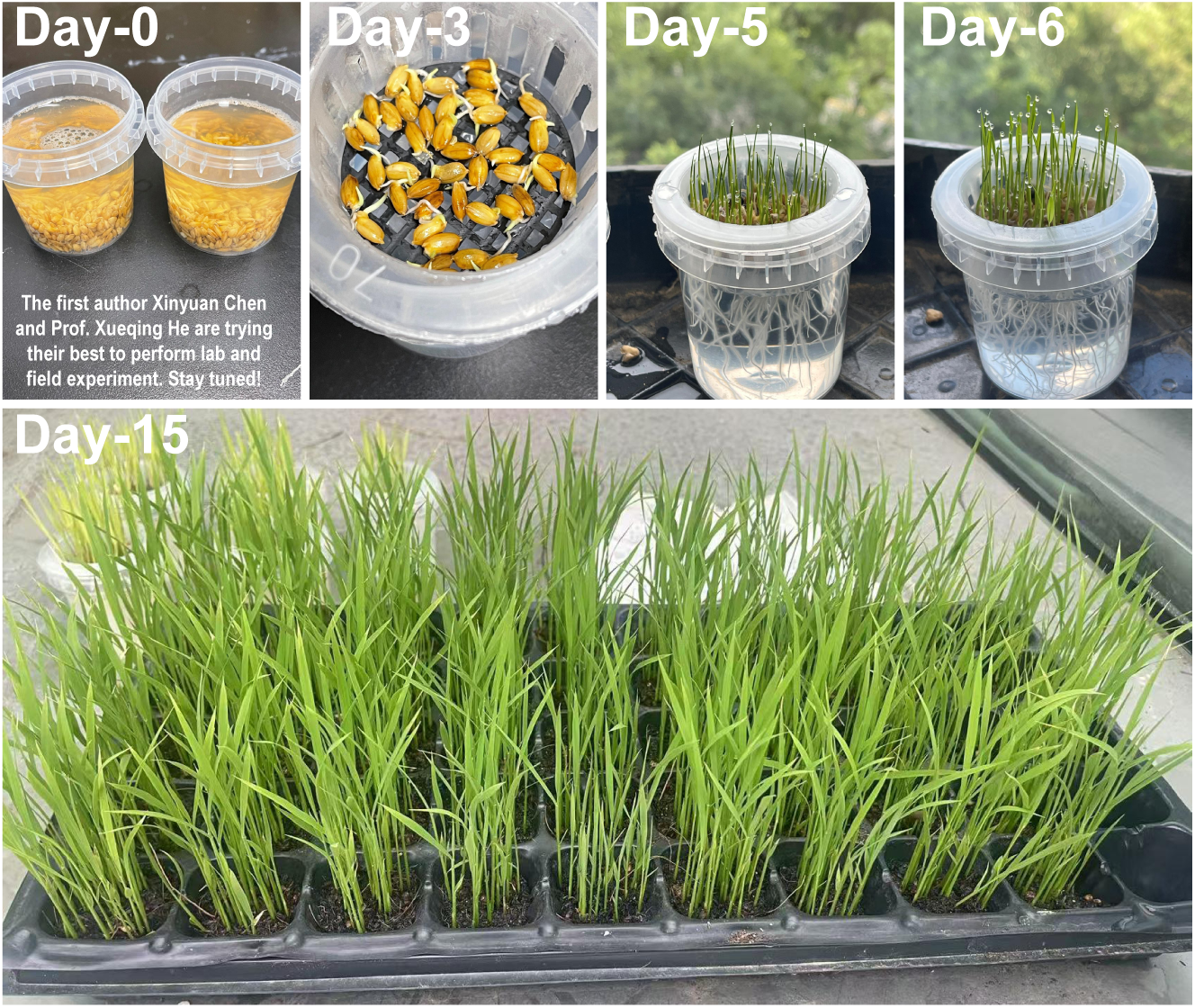
Growth of rice plants in lab. Those rice plants will be subjected to *in vitro* assays to determine lethal concentrations against BPH nymphs and adults.

Even all lab and field experimental results demonstrate promising efficacy of the FM and FW dipeptides against BPH, the practical implementation of this solution must carefully consider production costs to ensure economic viability for farmers. The dual motivations of this work, to replace hazardous chemical pesticides that endanger farmer/environment health while simultaneously avoiding solutions that would impose unsustainable financial burdens, remain central to our development strategy. Current preliminary cost analyses suggest these dipeptides could be economically competitive due to their simple molecular structure and potential for scalable synthesis, though detailed production optimization studies are needed to confirm this advantage. The true agricultural value proposition lies not just in the peptides’ biological activity, but in developing a cost-effective production and delivery system that maintains farmer profitability while addressing the health and environmental concerns of conventional pesticides.

Last but not the least, the biodegradable nature of peptide pesticides offers an exciting “one stone, two birds” opportunity, where these dipeptides (FM, FW) could serve dual roles as both effective BPH killers and valuable plant nutrient sources. When these peptides naturally decompose into amino acids like phenylalanine, methionine, and tryptophan, they may provide bioavailable amino acids or nitrogen and other essential nutrition elements that rice plants can utilize for growth, potentially creating a unique “pesticide-plus-fertilizer” effect. This potential dual functionality could transform sustainable agriculture by simultaneously addressing two major challenges: pest control and plant nutrition. The economic implications are profound - such multifunctional peptides could reduce farmers’ reliance on both chemical pesticides and synthetic fertilizers while improving crop yields. However, optimal application rates must be carefully determined to balance effective pest management with nutritional benefits, avoiding potential nutrition excess while maximizing the synergistic effects.

While substantial challenges lie ahead in scaling and optimizing this technology, our results demonstrate the exciting potential of multifunctional peptide pesticides. In summary, substantial work lies ahead to translate these results into real-world solutions.

## Conclusion

This study establishes a pioneering computational framework for designing next-generation peptide biopesticides targeting the n1AChR-*α*3 in the devastating rice pest BPH or *Nilaparvata lugens*. By integrating AF3-predicted structures with multi-force-field MD simulations, we resolved the atomic 3D structure of nAChR-*α*3 and identified conformational dynamics critical for ligand binding. Our innovative “Dipeptide Probing” strategy - systematically screening 20 phenylalanine-containing dipeptides, revealed FM (Phe-Met) as the highest-affinity binder, stabilized by AIHB mechanism, where FM dipeptides self-assemble via HB to form stable, protein target-engaged clusters through hydrophobic interactions, challenging reductionist views of peptide-receptor interactions. Beyond identifying FM as a lead biopesticide candidate, this work provides a high-resolution structural blueprint for rational pesticide design, a generalizable methodology for mapping protein binding sites, and a sustainable path for agriculture through biodegradable peptides that minimize ecological harm while potentially acting as nutrient sources upon degradation. Future efforts will validate FM/FW efficacy in lab and field trials, assess ecotoxicological impacts, and optimize cost-effective production, collectively advancing a MD-guided pipeline that transforms structure-based agrochemical discovery toward eco-friendly pest management solutions.

## Acknowledgement

Dr. Jiaqi Wang acknowledges the funding support of the National Natural Science Foundation of China (No. 52101023) and Basic Research Program of Jiangsu - General Program of Jiangsu Provincial Department of Science and Technology (No. BK20241816). Dr. Xueqing He acknowledges the funding support of Research Development Fund (RDF) of Xi’an Jiaotong-Liverpool University (RDF-22-02-043). The authors also acknowledge the high-performance computing platform at Xi’an Jiaotong-Liverpool University and Beijing Paratera Tech Corp. LTD.

## Supporting Information Available

RMSD evolutions of 400 dipeptides from 0 to 500 ns in MD simulations.

## Extended Data of Dipeptide-nAChR-***α***3 Interactions

**Special note 1:** Peptides shown in Extended Data Figure 1 to 16 were all screened based on conformational stability, quantified by time-averaged RMSD values calculated over the equilibrated trajectory window (300–500 ns), different from the STD evaluations as shown in Figure 5c. Candidates exhibiting an average RMSD *<* 0.5 nm relative to the protein receptor binding site were prioritized as stable binders here.

**Special note 2 (color scheme):** In Extended Data Figures 1–16, backbone carbon atoms of dipeptides are consistently colored **yellow**. In contrast, backbone carbon atoms in the protein are primarily **purple** (flexible loops) and **forest green** (*α*-helices), with a small subset appearing **navy blue** (*β*-sheets). Nitrogen and oxygen atoms retain standard coloring (**blue** and **red**, respectively) in both the protein and dipeptides.

**Extended Data Figure 1:**
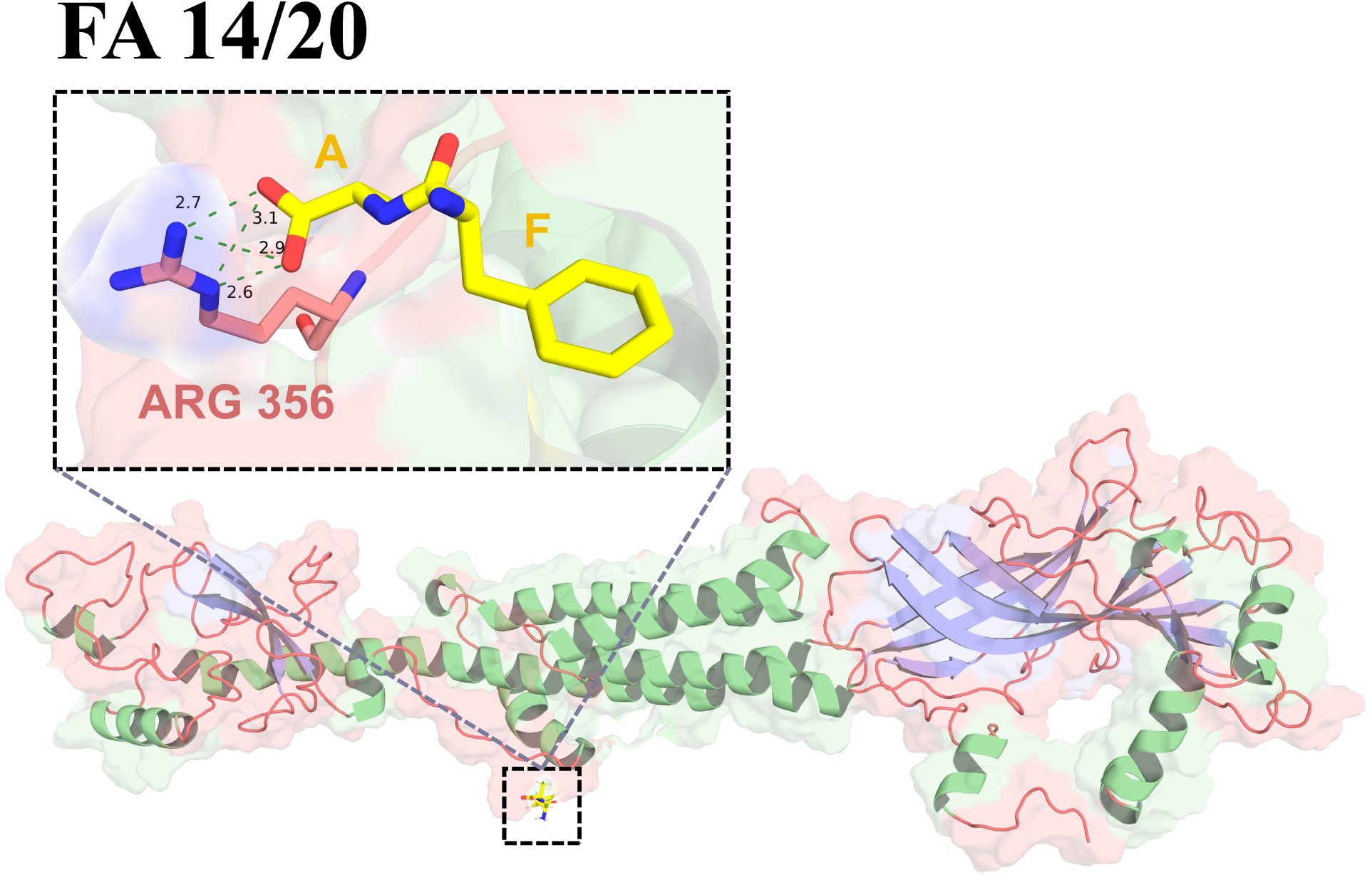
PyMOL-based Interaction analysis of the **FA**-nAChR-*α*3 complex identifies **ARG 356** as a critical binding residue, with persistent HBs (indicated by dashes with distances) observed throughout the equilibrated trajectory from 300 to 500 ns.

**Extended Data Figure 2:**
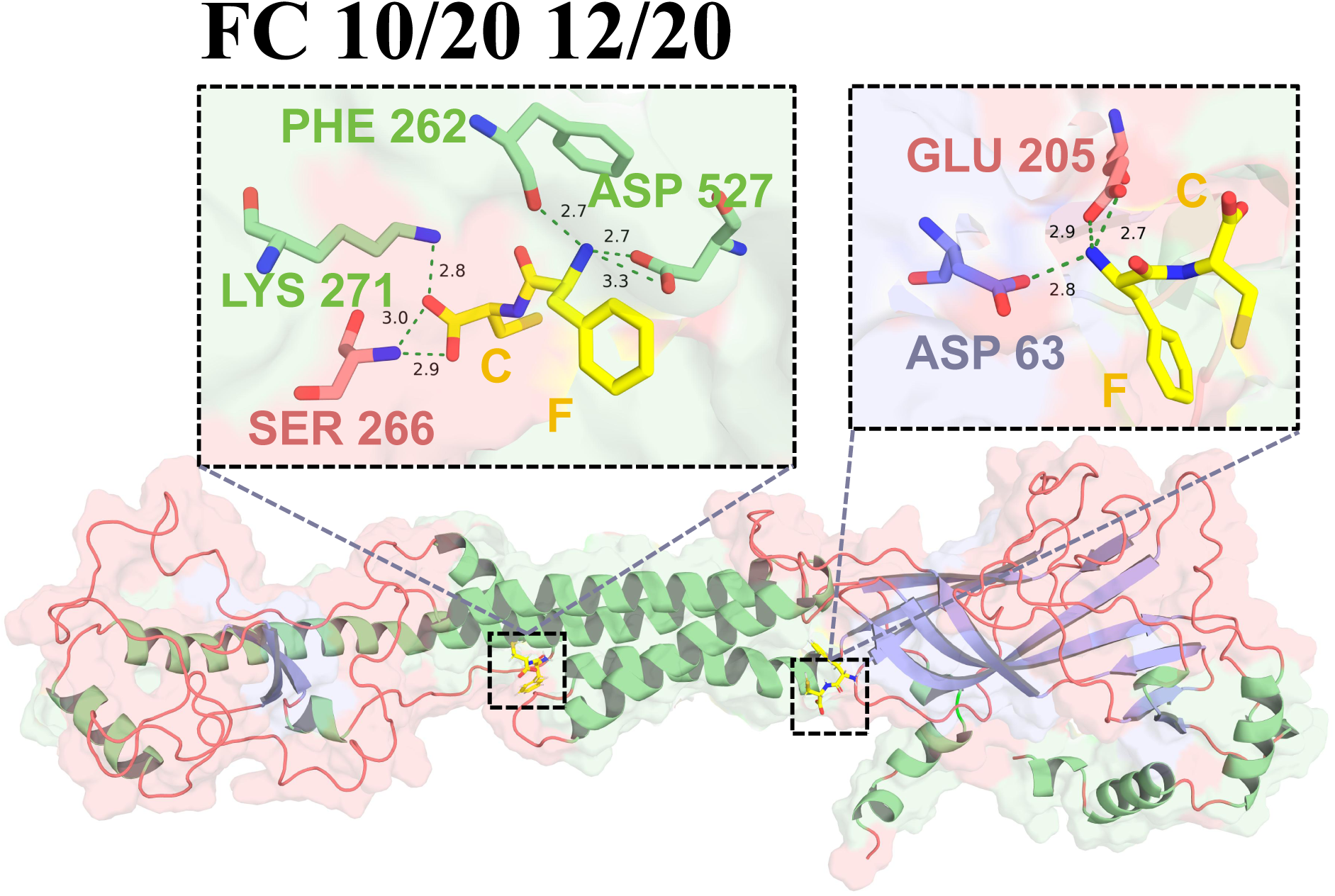
Interaction analysis of the **FC**-nAChR-*α*3 complex. The FC locates at two sites, and interacts with residueS **SER 266**, **LYS 271**, **PHE 262**, and **ASP 527** at one site, and interacts with **ASP 63** and **GLU 205** at another site.

**Extended Data Figure 3:**
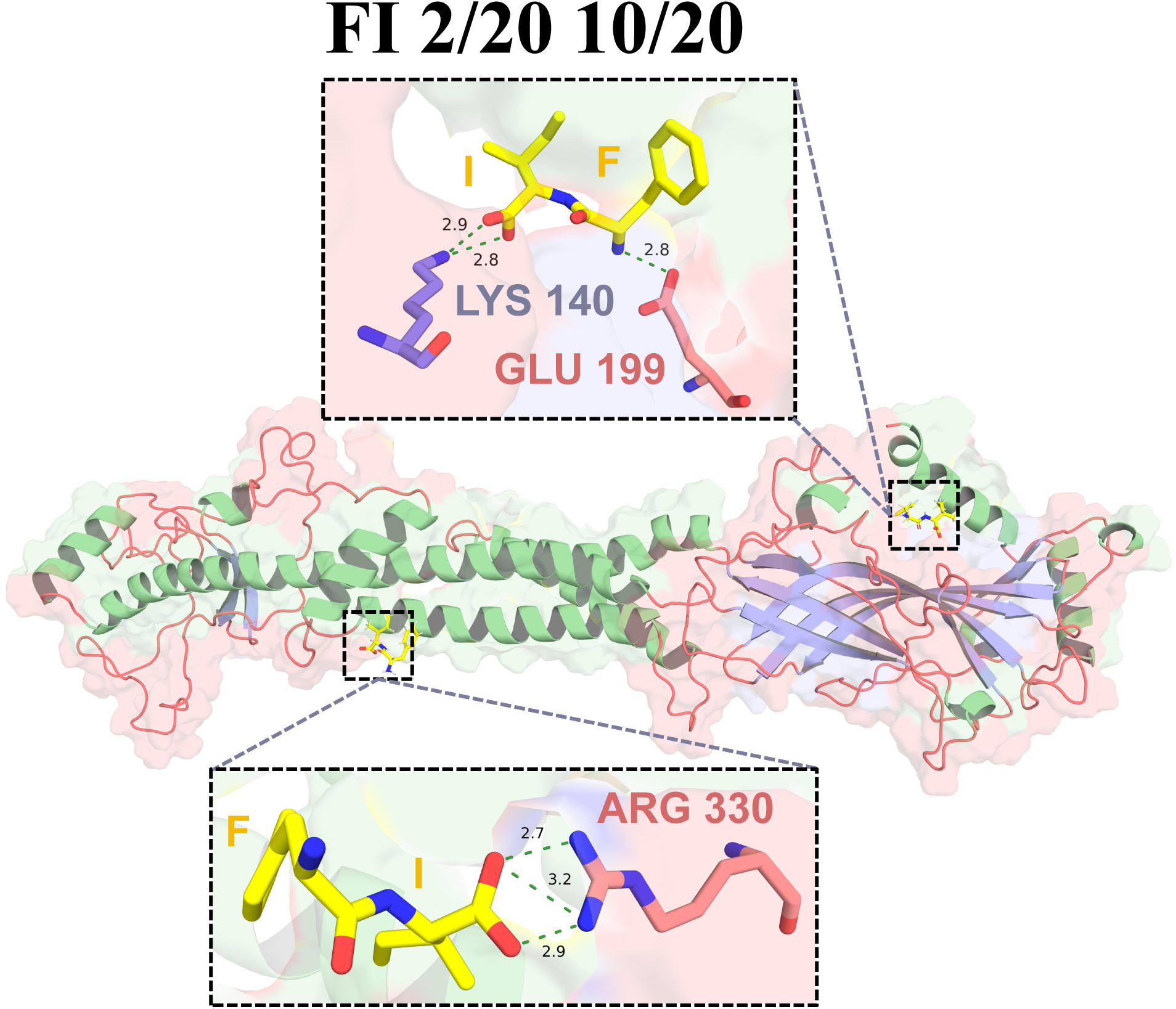
Interaction analysis of the **FI**-nAChR-*α*3 complex. The FI locates at two sites, and interacts with residue **LYS 140** and **GLU 199** at one site, and interacts with **ARG 330** at another site.

**Extended Data Figure 4:**
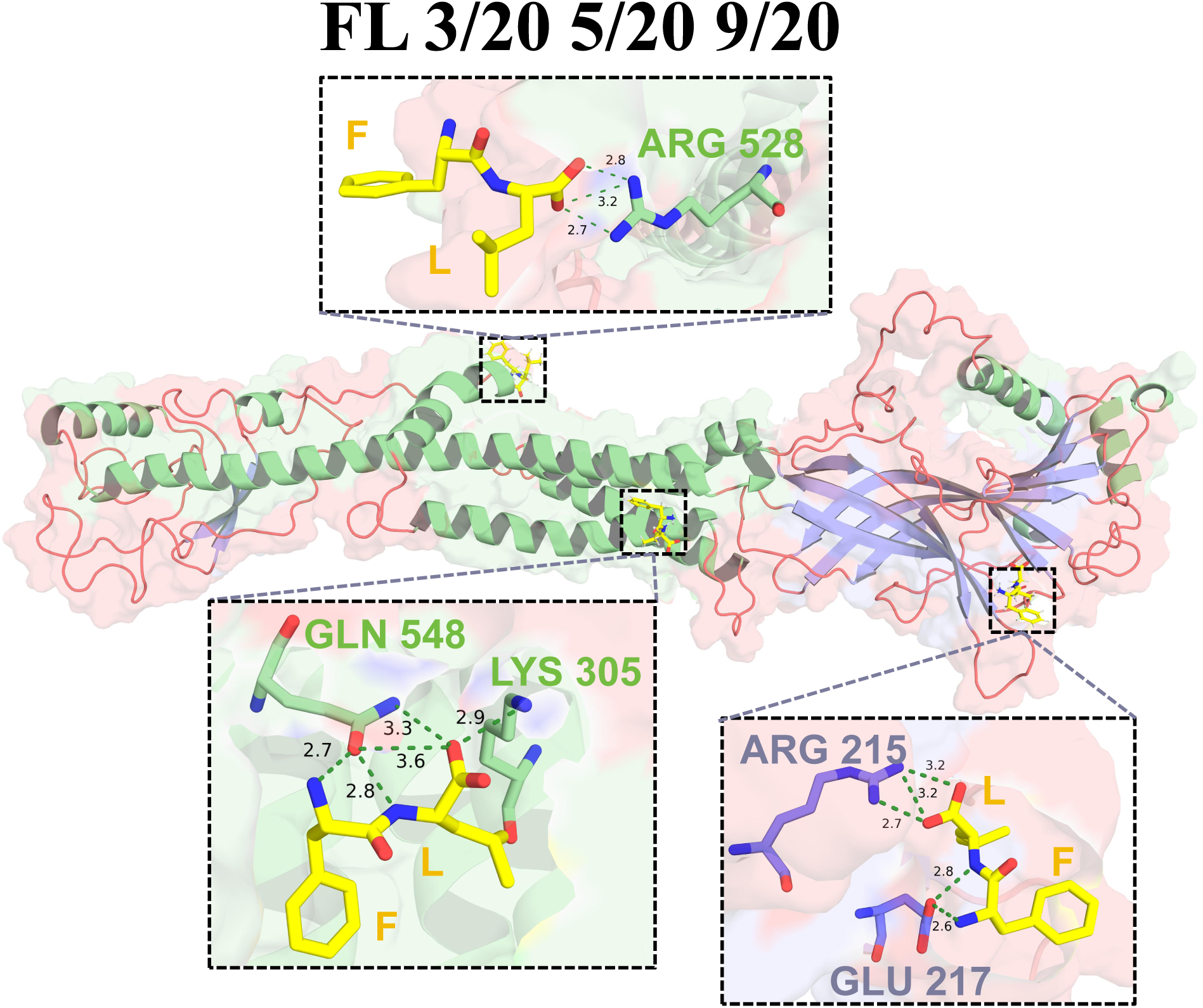
Interaction analysis of the **FL**-nAChR-*α*3 complex. The FL locates at three sites, and interacts with residue **ARG 528** at the first site, **GLN 548** and **LYS 305** at the second site, and **GLU 217** and **ARG 215** at the third site.

**Extended Data Figure 5:**
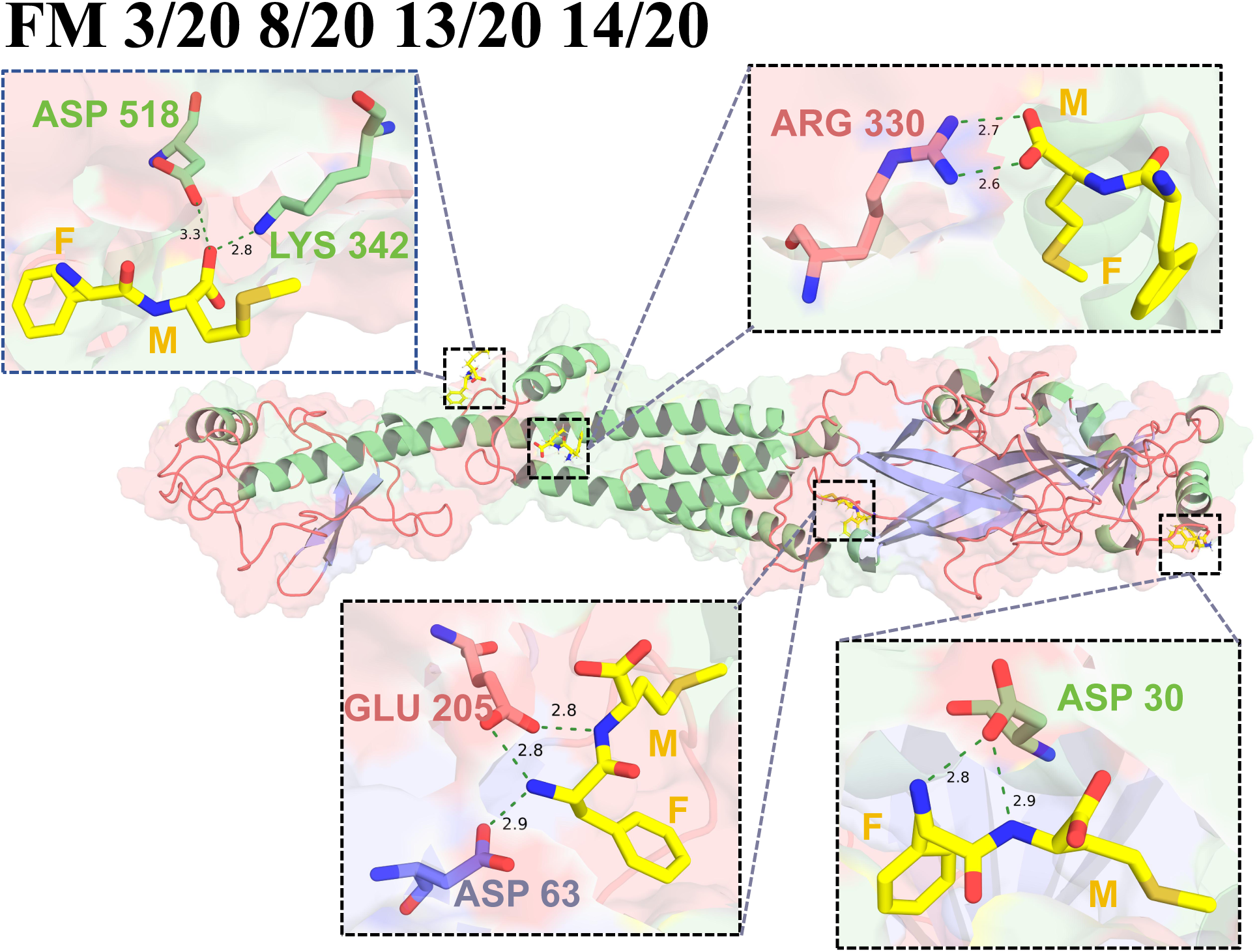
Interaction analysis of the **FM**-nAChR-*α*3 complex. The FM locates at four sites, and interacts with residue **ASP 518** and **LYS 342** at the first site (also illustrated as A2 in Figure 7), with **ARG 330** at the second site (A3 in Figure 7), with **GLU 205** and **ASP 63** at the third site (A4 in Figure 7), and interacts with **ASP 30** at the fourth site (A5 in Figure 7). However, through the RMSD analysis, the interactions of FM with residues ARG 481, ARG 482, and HIS 485 (shown as A1 in Figure 7) are missing here.

**Extended Data Figure 6:**
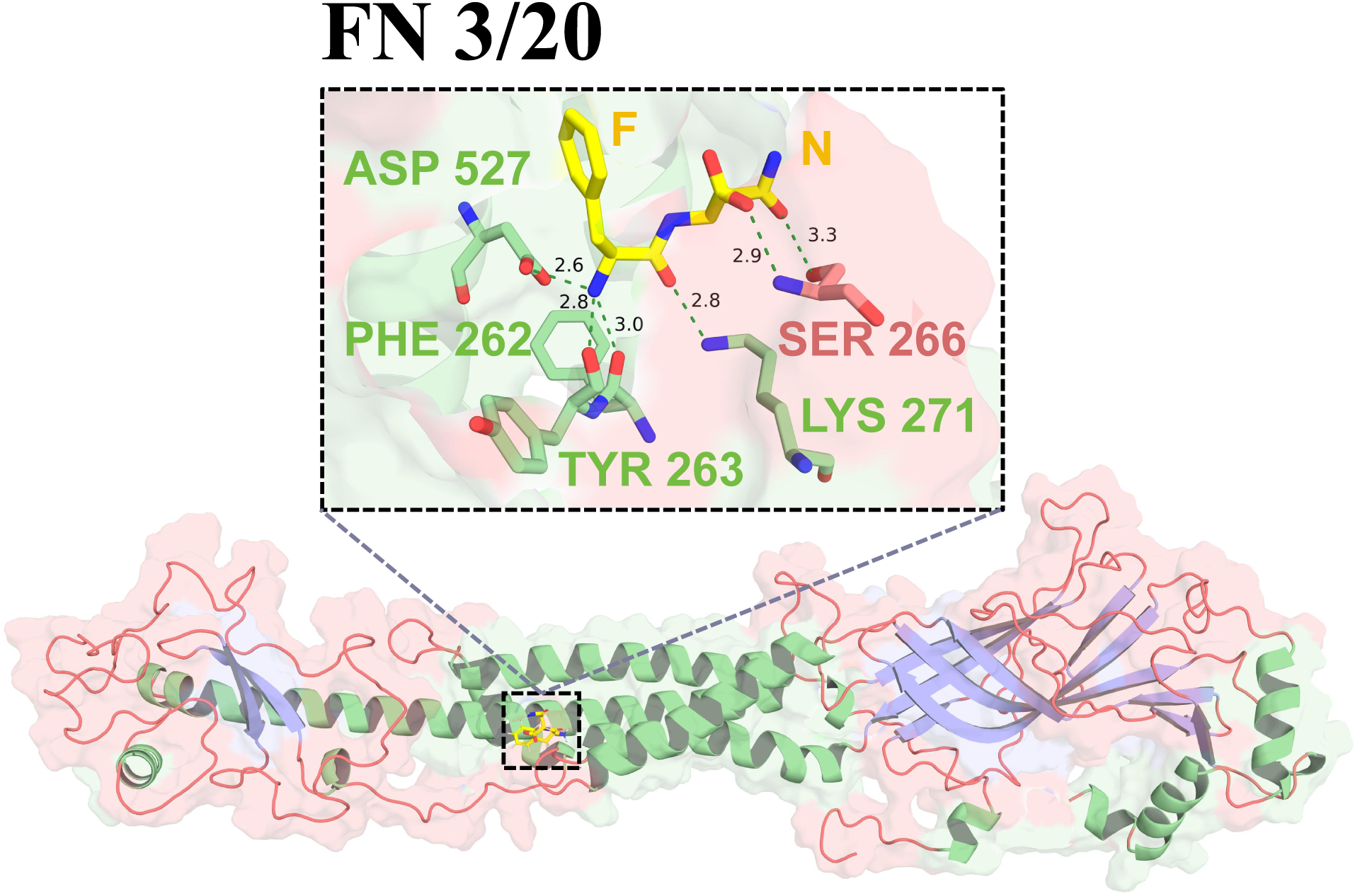
Interaction analysis of the **FN**-nAChR-*α*3 complex. The FN locates at only one site, and but interacts with five residues **SER 266**, **LYS 271**, **TYR 263**, **PHE 262** and **ASP 527**.

**Extended Data Figure 7:**
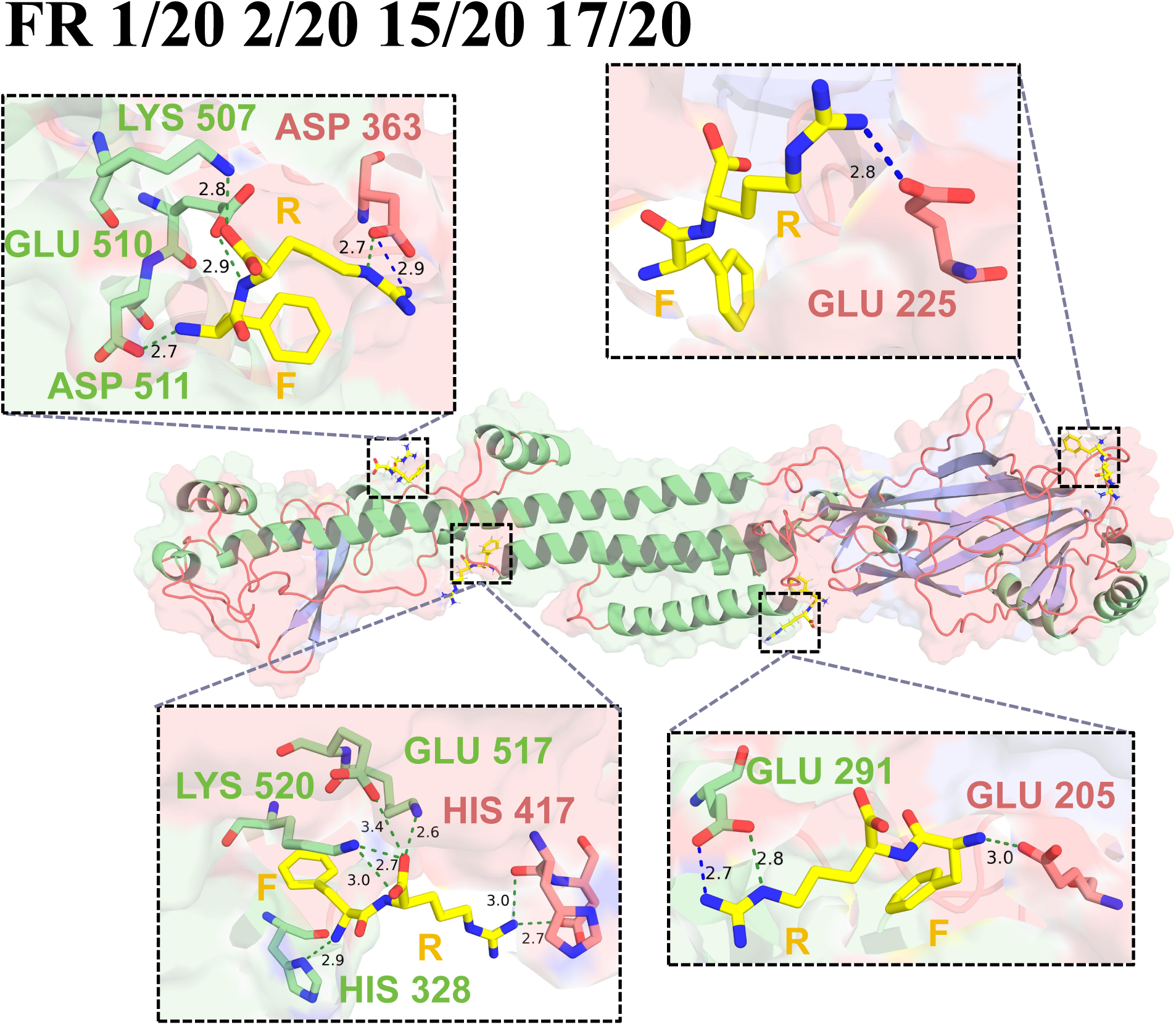
Interaction analysis of the **FR**-nAChR-*α*3 complex. The FR locates at four sites, and interacts with residues **LYS 507**, **GLU 510**, **ASP 511**, and **ASP 363** at the first site, with residue **GLU 225** at the second site, with residues **LYS 520**, **GLU 517**, **HIS 417**, and **HIS 328** at the third site, and interacts with residues **GLU 291** and **GLU 205** at the fourth site.

**Extended Data Figure 8:**
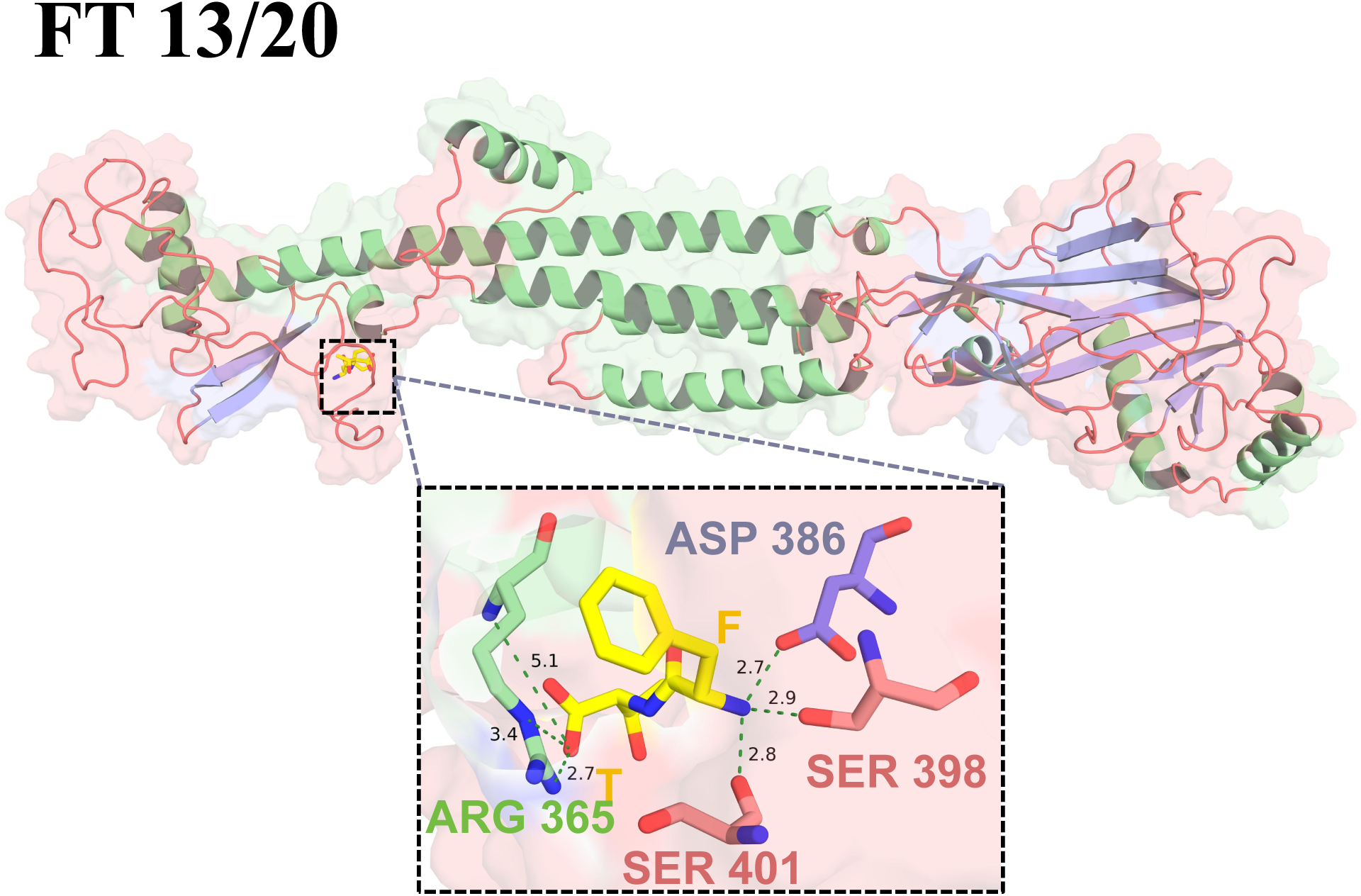
Interaction analysis of the **FT**-nAChR-*α*3 complex. The FT locates at only one site, and interacts with four residues, i.e., **ARG 365**, **SER 401**, **SER 398**, and **ASP 386**.

**Extended Data Figure 9:**
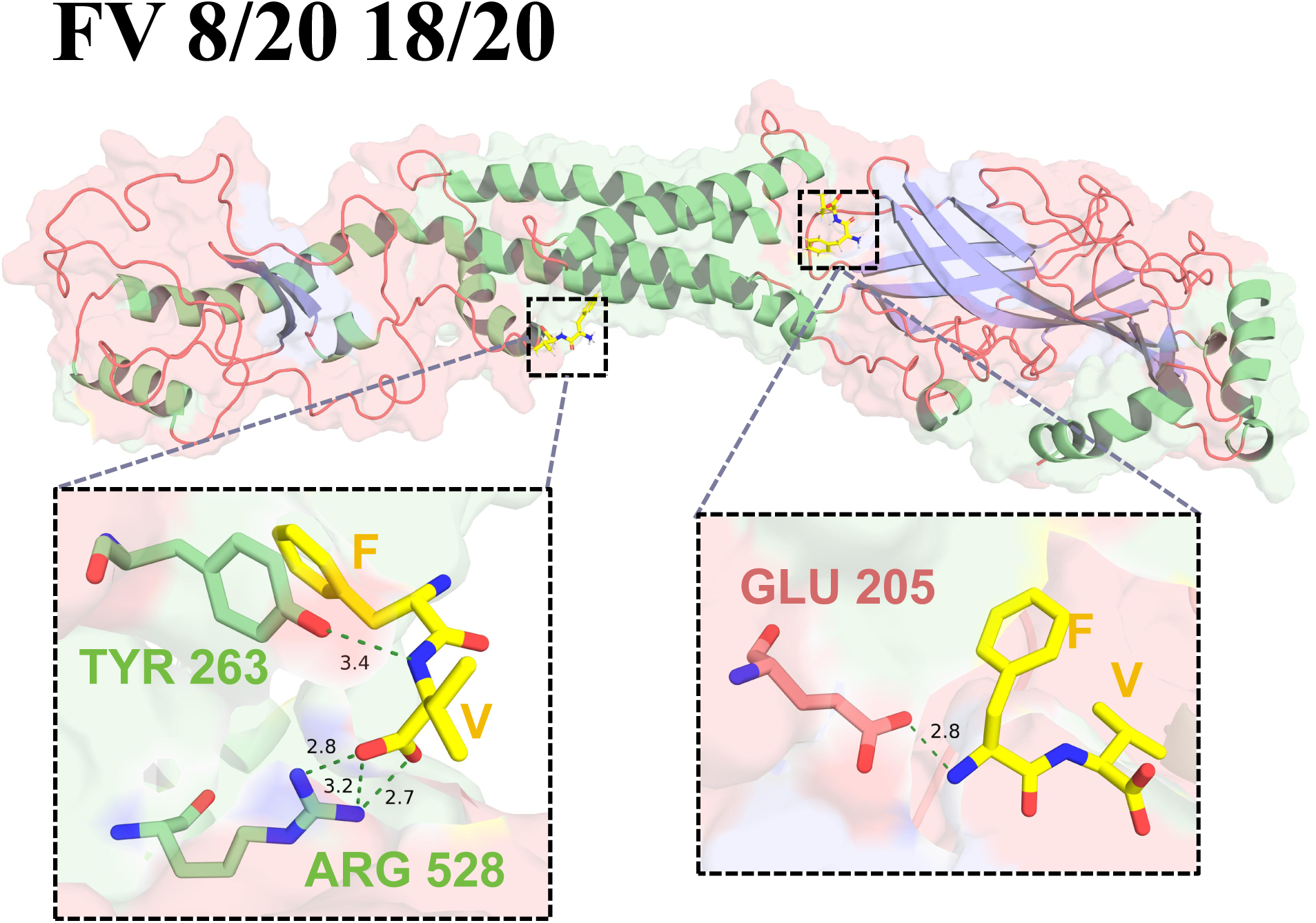
Interaction analysis of the **FV**-nAChR-*α*3 complex. The FV locates at two sites, and interacts with residues **TYR 263** and **ARG 528** at the first site, and interacts with residue **GLU 205** at the second site.

**Extended Data Figure 10:**
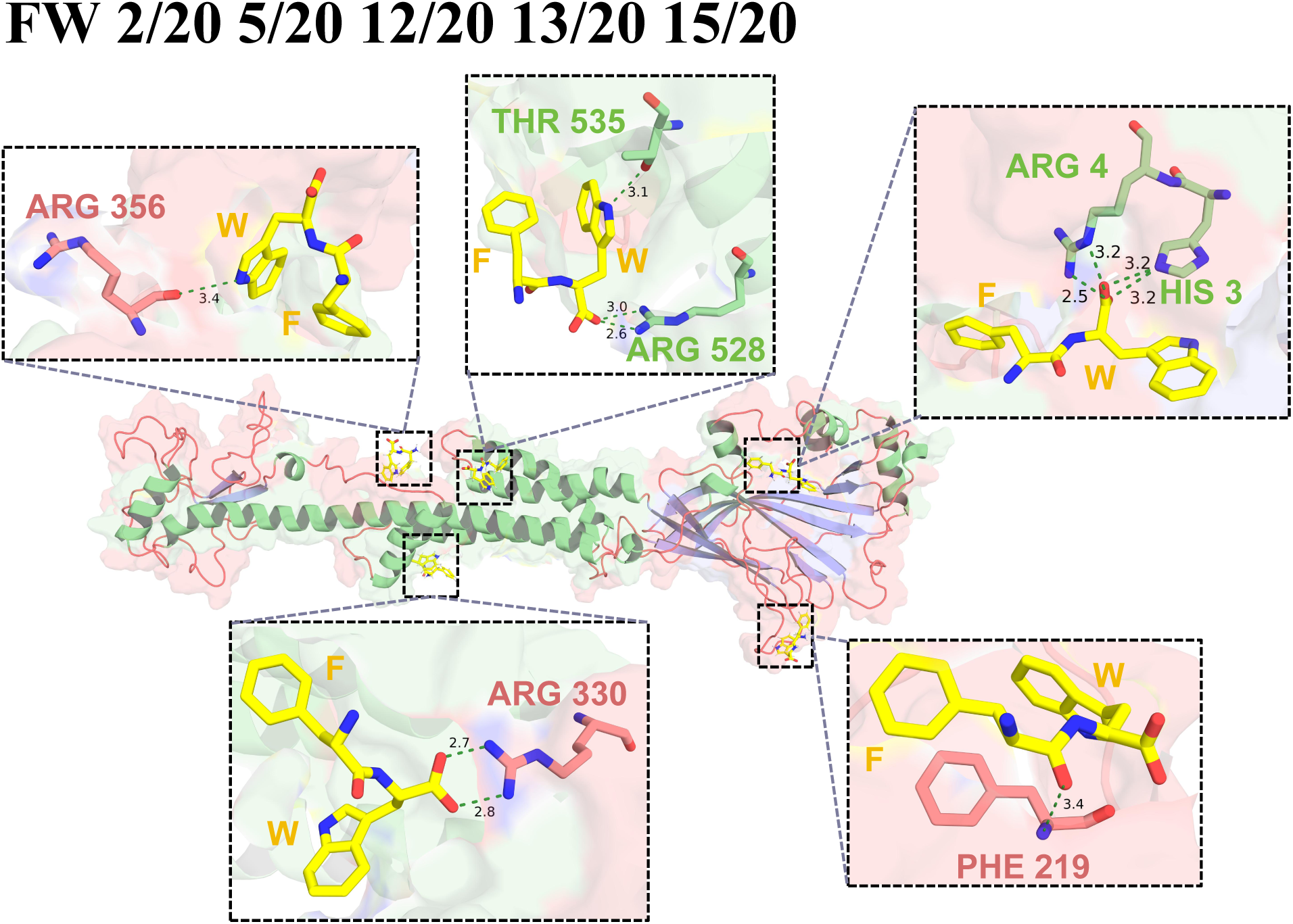
Interaction analysis of the **FW**-nAChR-*α*3 complex. The FW locates at five sites, and interacts with residue **ARG 356** at the first site, with residues **THR 535** and **ARG 528** at the second site, with residues **ARG 4** and **HIS 3** at the third site, with residue **ARG 330** at the fourth site, and interacts with residue **PHE 219** at the fifth site.

**Extended Data Figure 11:**
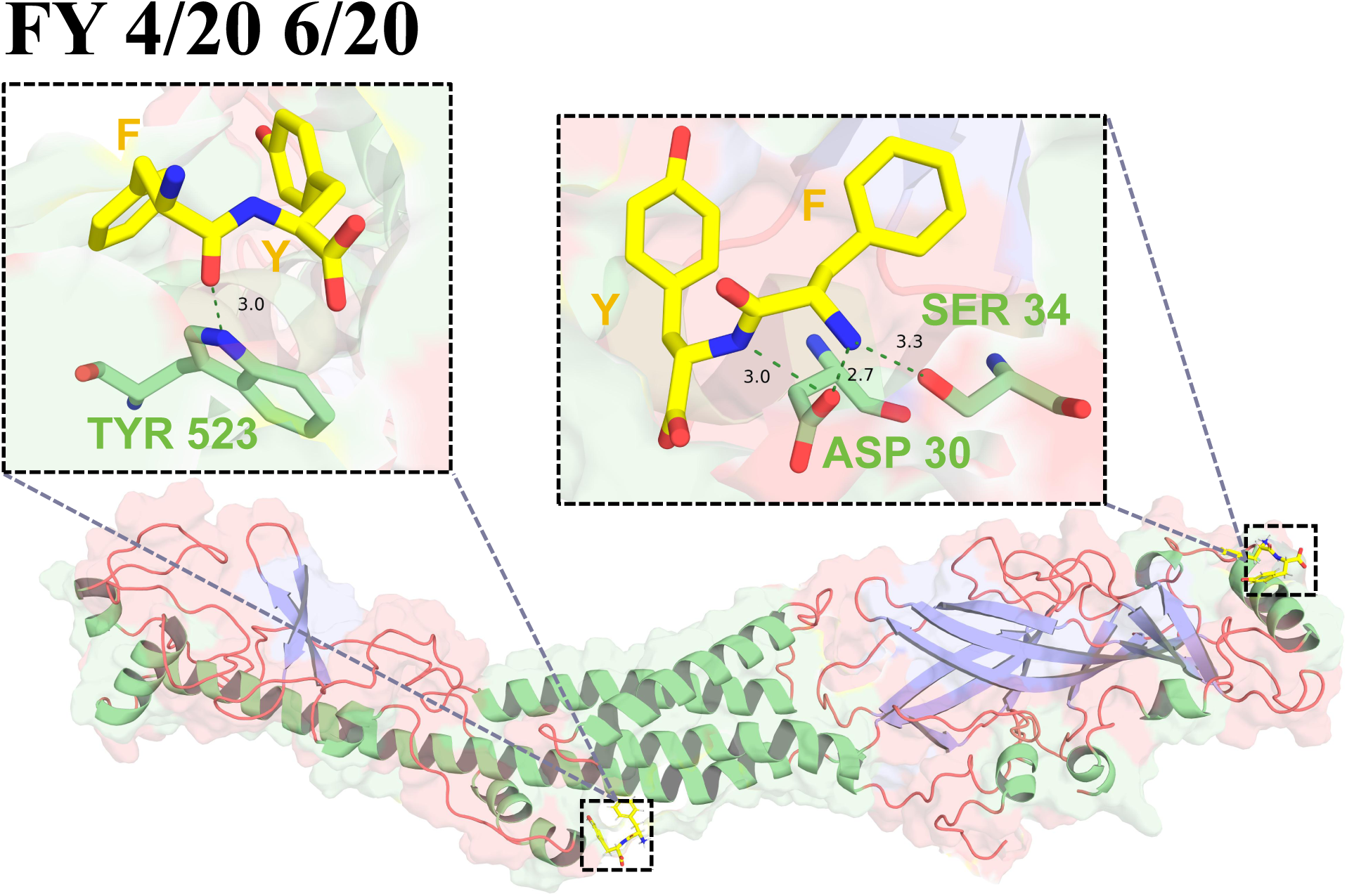
Interaction analysis of the **FY**-nAChR-*α*3 complex. The FY locates at two sites, and interacts with residue **TYR 523** at the first site, and interacts with residues **ASP 30** and **SER 34** at the second site.

**Extended Data Figure 12:**
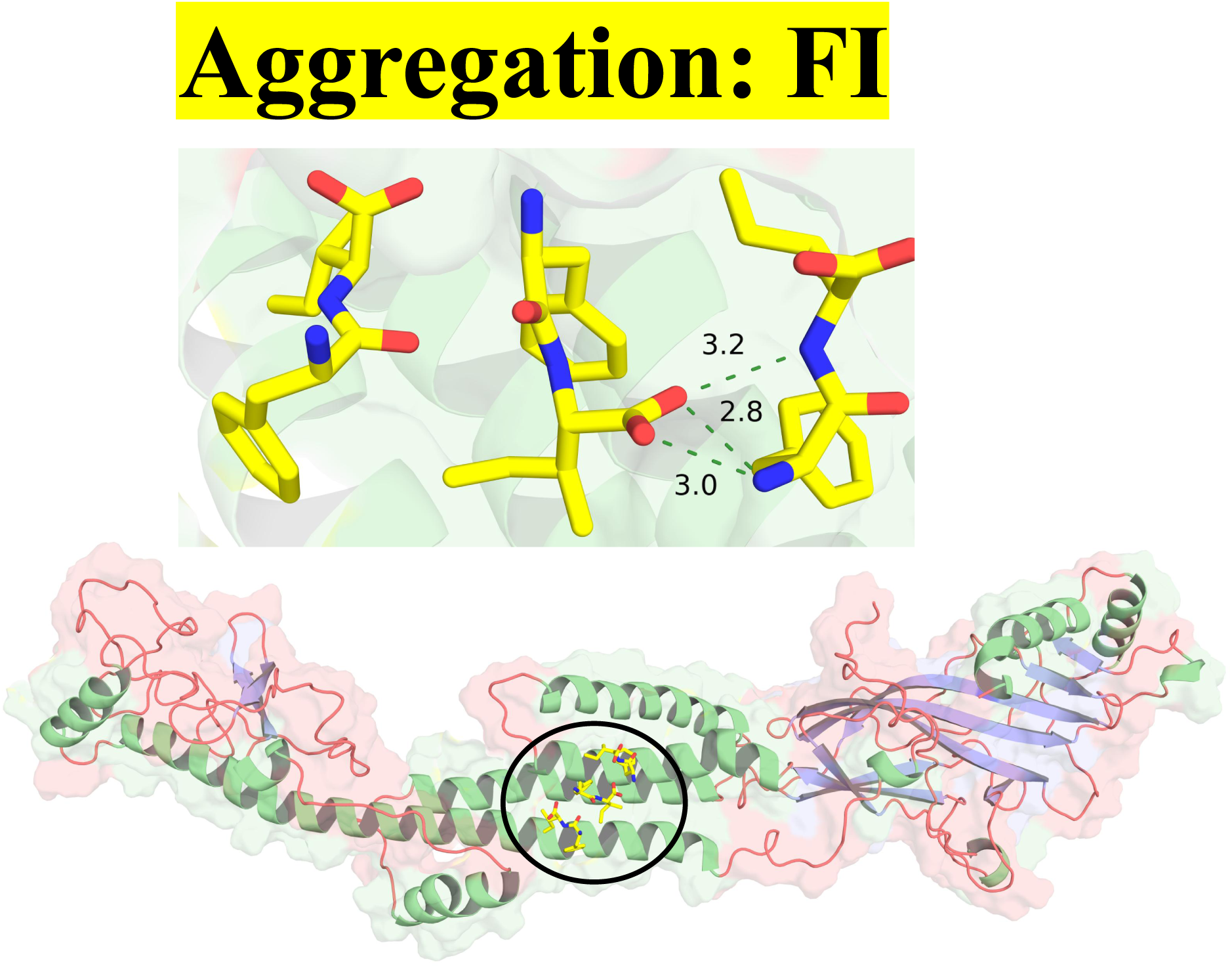
Aggregation within FI dipeptides.

**Extended Data Figure 13:**
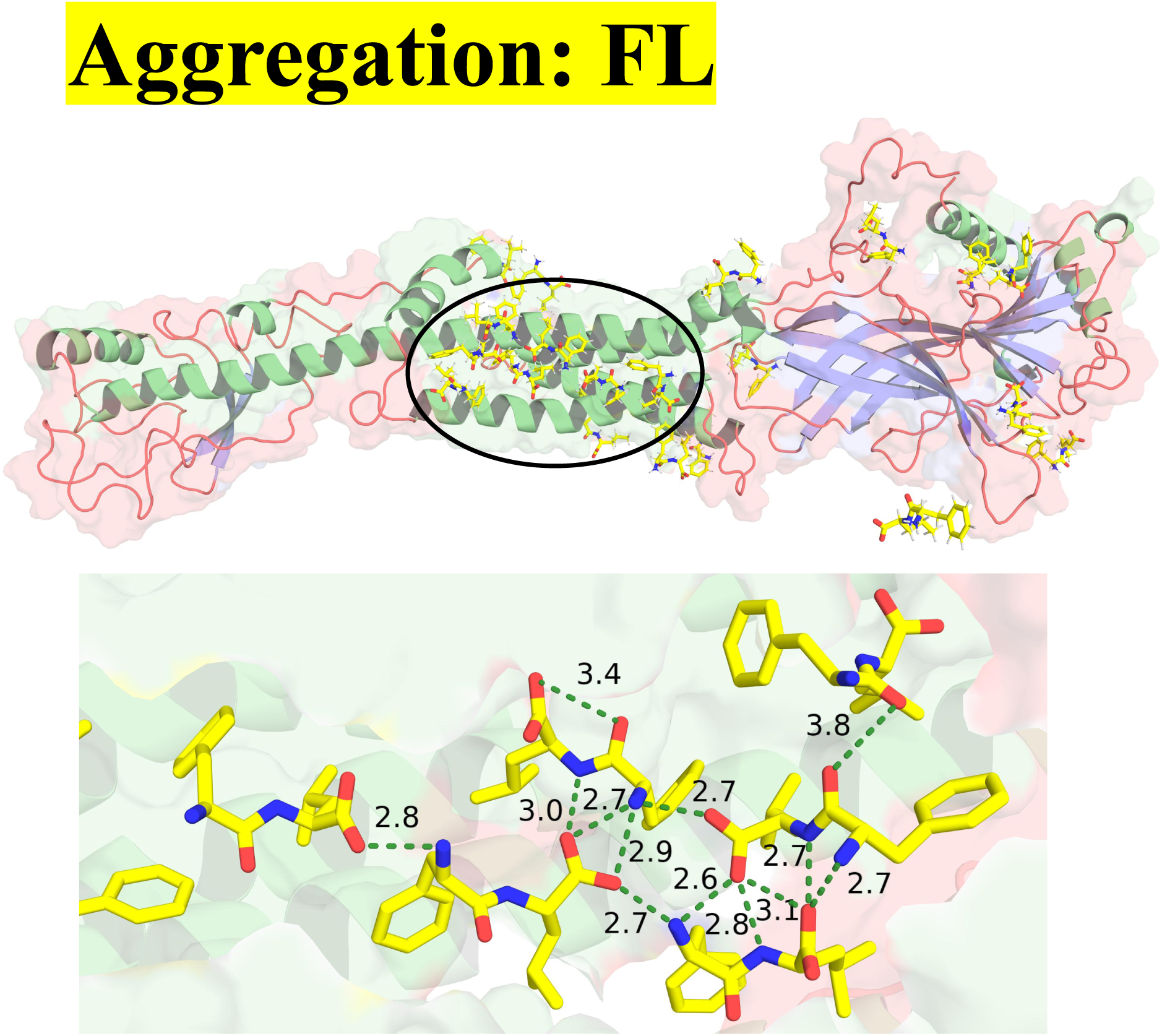
Aggregation within FL dipeptides.

**Extended Data Figure 14:**
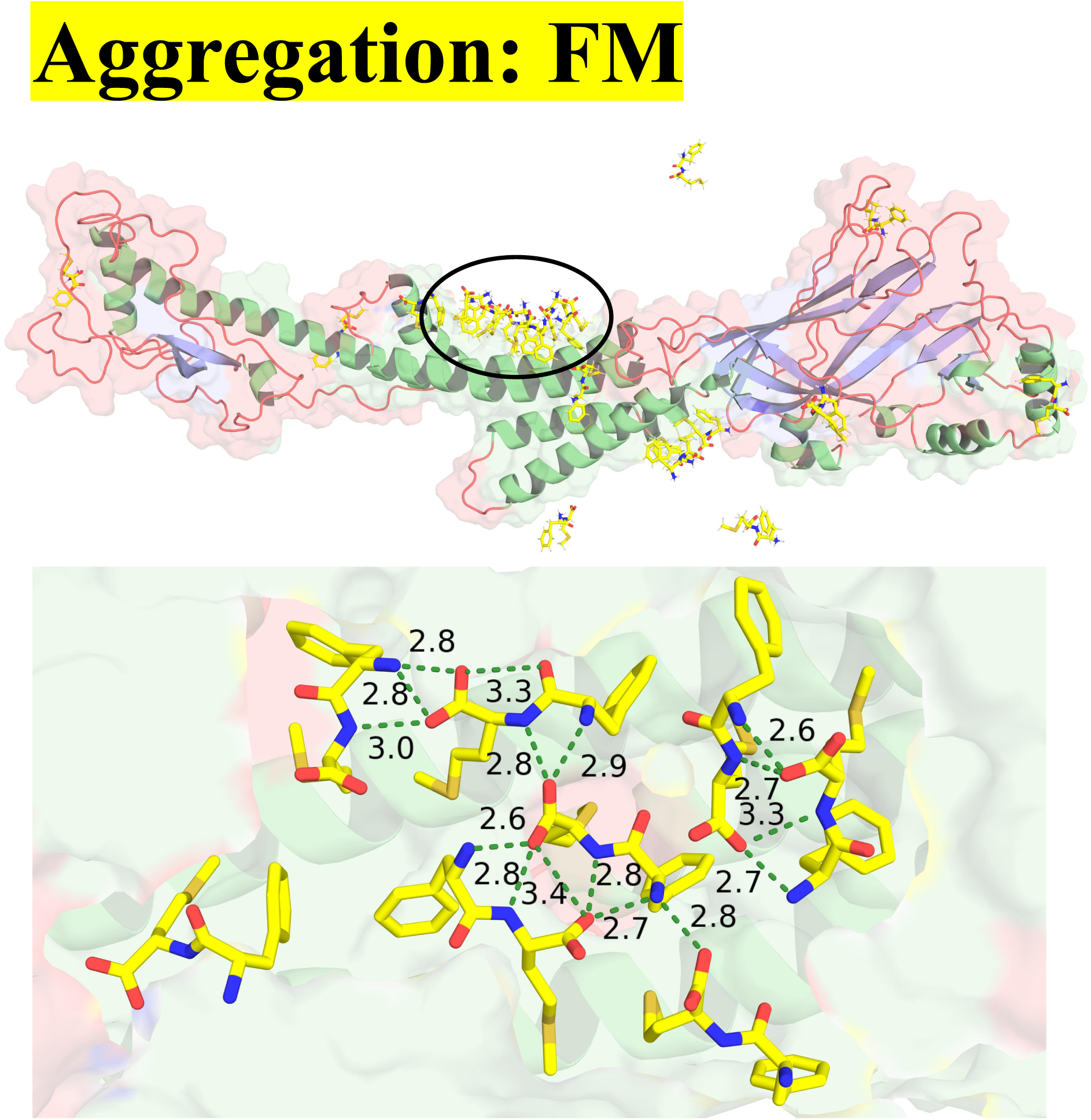
Aggregation within FM dipeptides (same as in Figure 6, last panel).

**Extended Data Figure 15:**
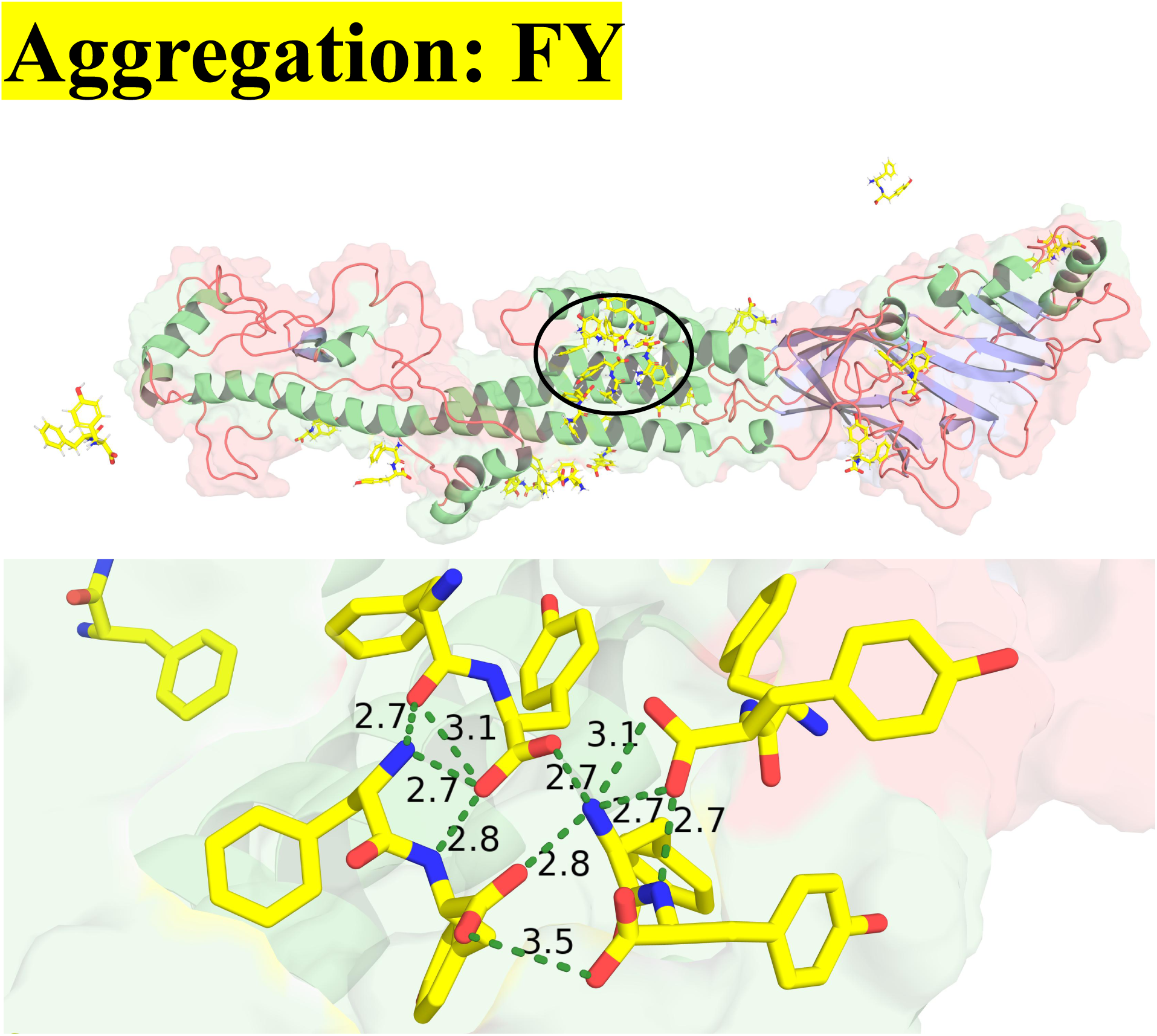
Aggregation within FY dipeptides.

**Extended Data Figure 16:**
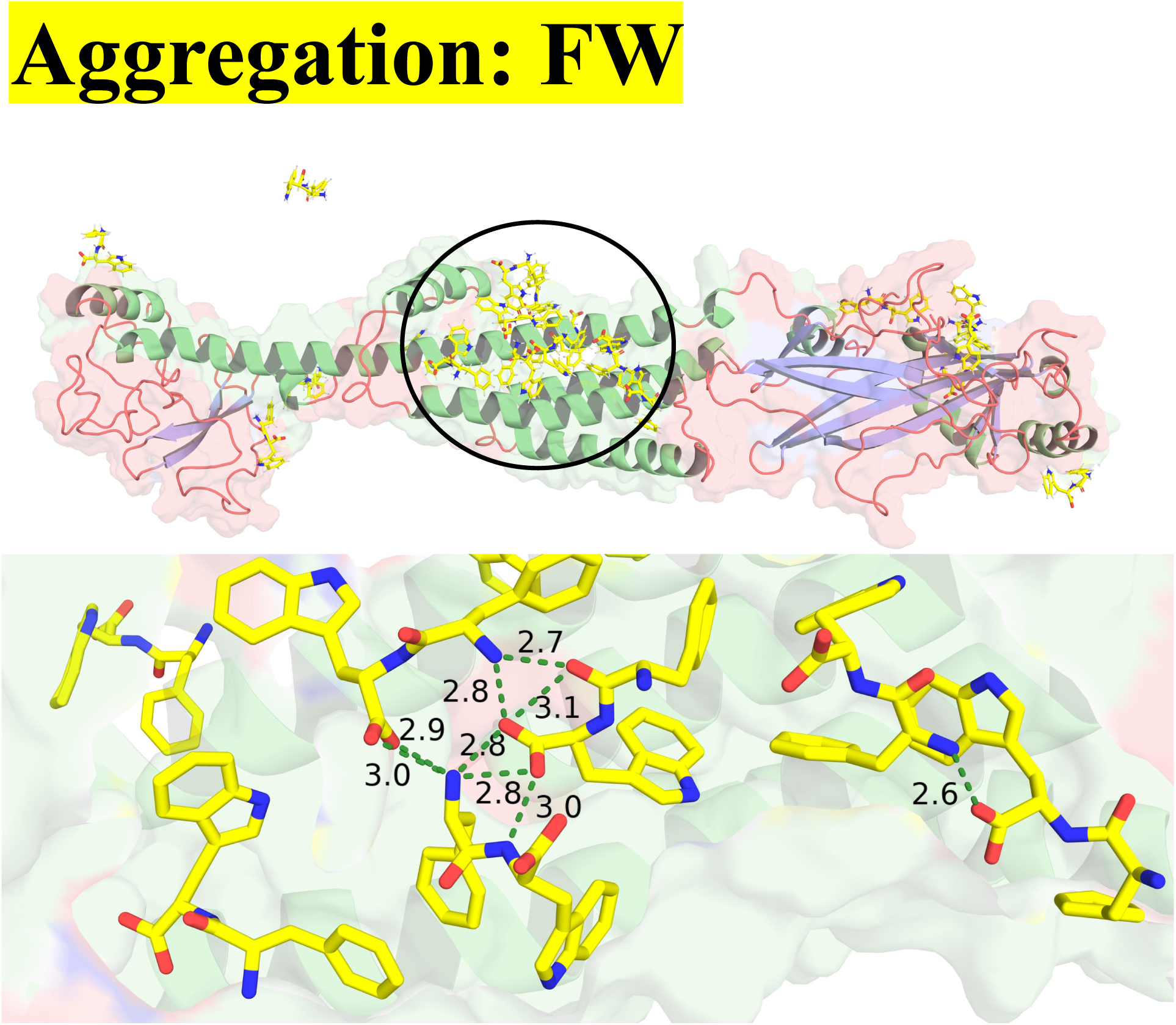
Aggregation within FW dipeptides.

