## Supplementary Data for "Computational Design of Next-Gen Peptide Biopesticides: Targeting the Nicotinic Acetylcholine Receptor in Rice Pests"

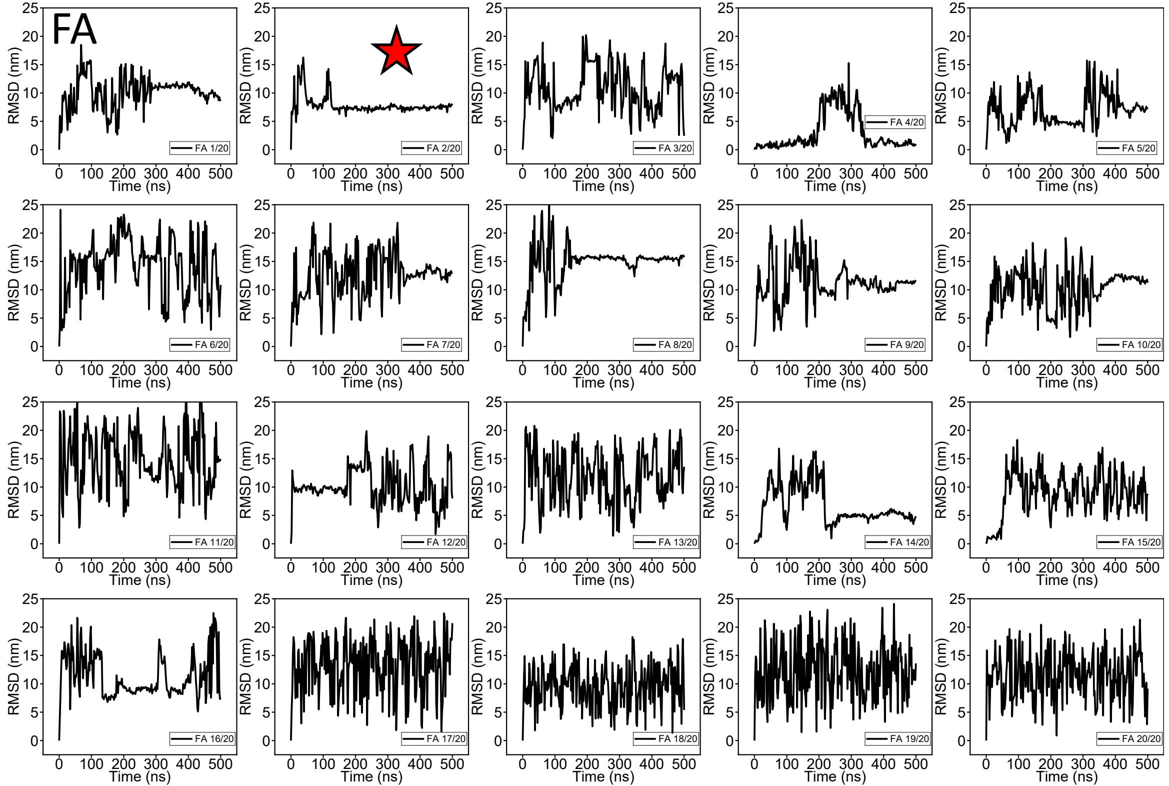
FA
RMSD (nm)
Time (ns)
FA 1/20
FA 2/20
FA 3/20
FA 4/20
FA 5/20
FA 6/20
FA 7/20
FA 8/20
FA 9/20
FA 10/20
FA 11/20
FA 12/20
FA 13/20
FA 14/20
FA 15/20
FA 16/20
FA 17/20
FA 18/20
FA 19/20
FA 20/20

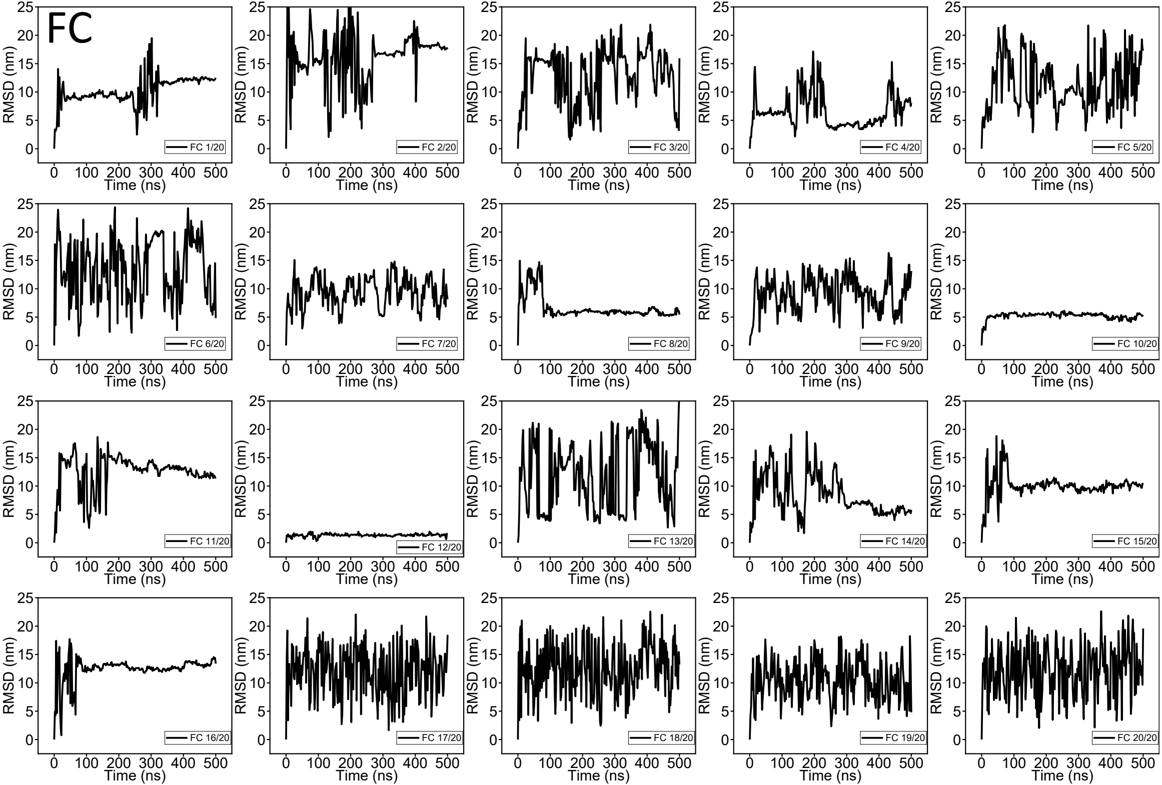
FC
RMSD (nm)
Time (ns)
FC 1/20
FC 2/20
FC 3/20
FC 4/20
FC 5/20
FC 6/20
FC 7/20
FC 8/20
FC 9/20
FC 10/20
FC 11/20
FC 12/20
FC 13/20
FC 14/20
FC 15/20
FC 16/20
FC 17/20
FC 18/20
FC 19/20
FC 20/20

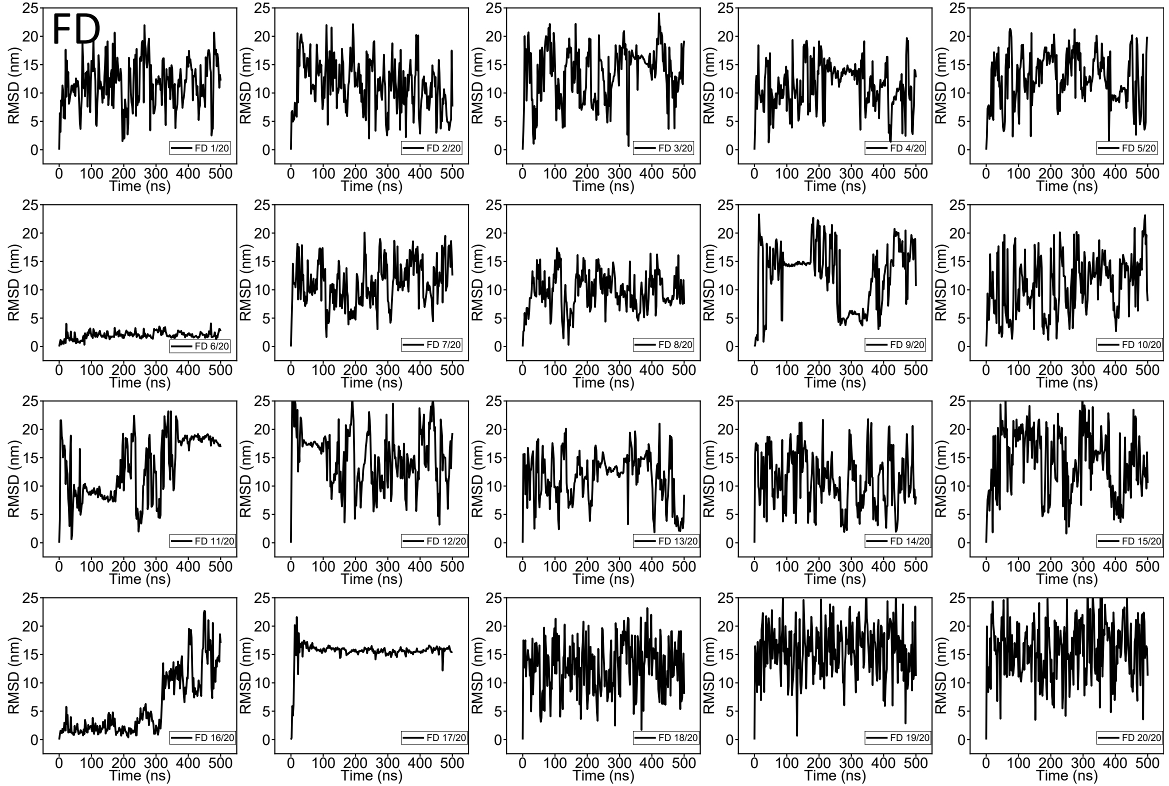

FD
RMSD (nm)
Time (ns)
FD 1/20
FD 2/20
FD 3/20
FD 4/20
FD 5/20
FD 6/20
FD 7/20
FD 8/20
FD 9/20
FD 10/20
FD 11/20
FD 12/20
FD 13/20
FD 14/20
FD 15/20
FD 16/20
FD 17/20
FD 18/20
FD 19/20
FD 20/20

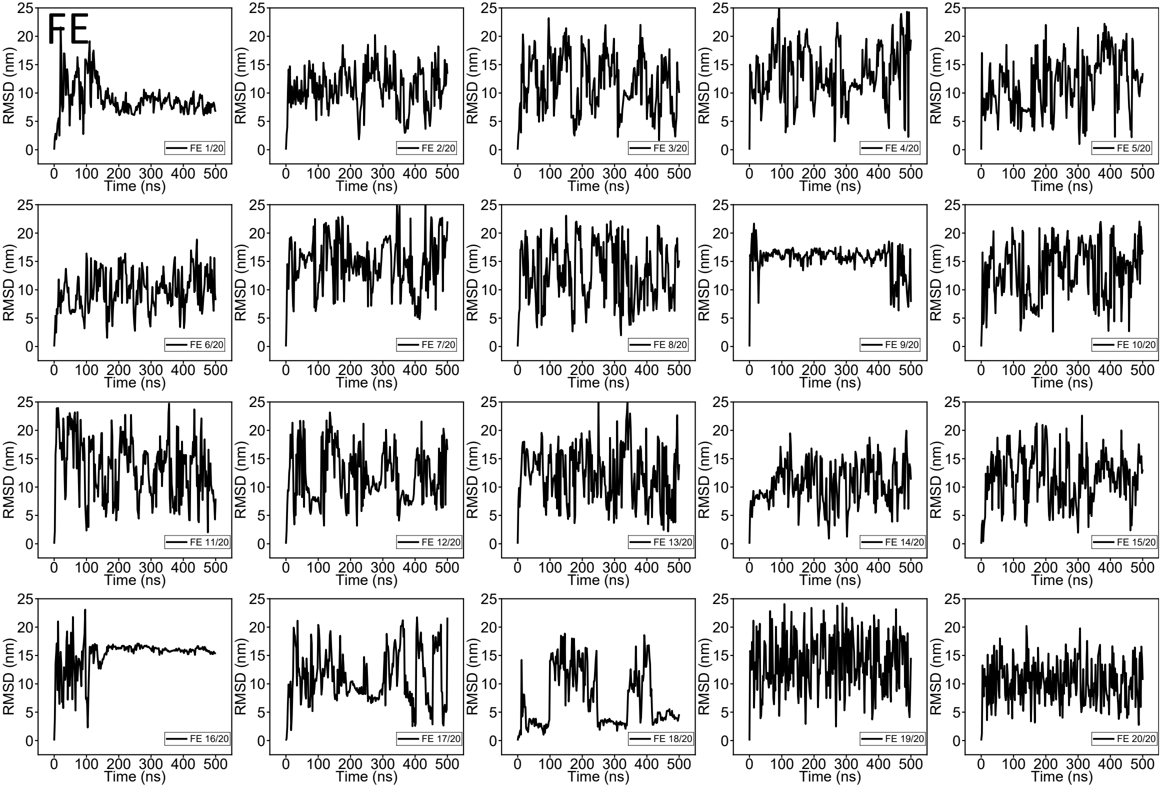
FE
RMSD (nm)
Time (ns)
FE 1/20
FE 2/20
FE 3/20
FE 4/20
FE 5/20
FE 6/20
FE 7/20
FE 8/20
FE 9/20
FE 10/20
FE 11/20
FE 12/20
FE 13/20
FE 14/20
FE 15/20
FE 16/20
FE 17/20
FE 18/20
FE 19/20
FE 20/20

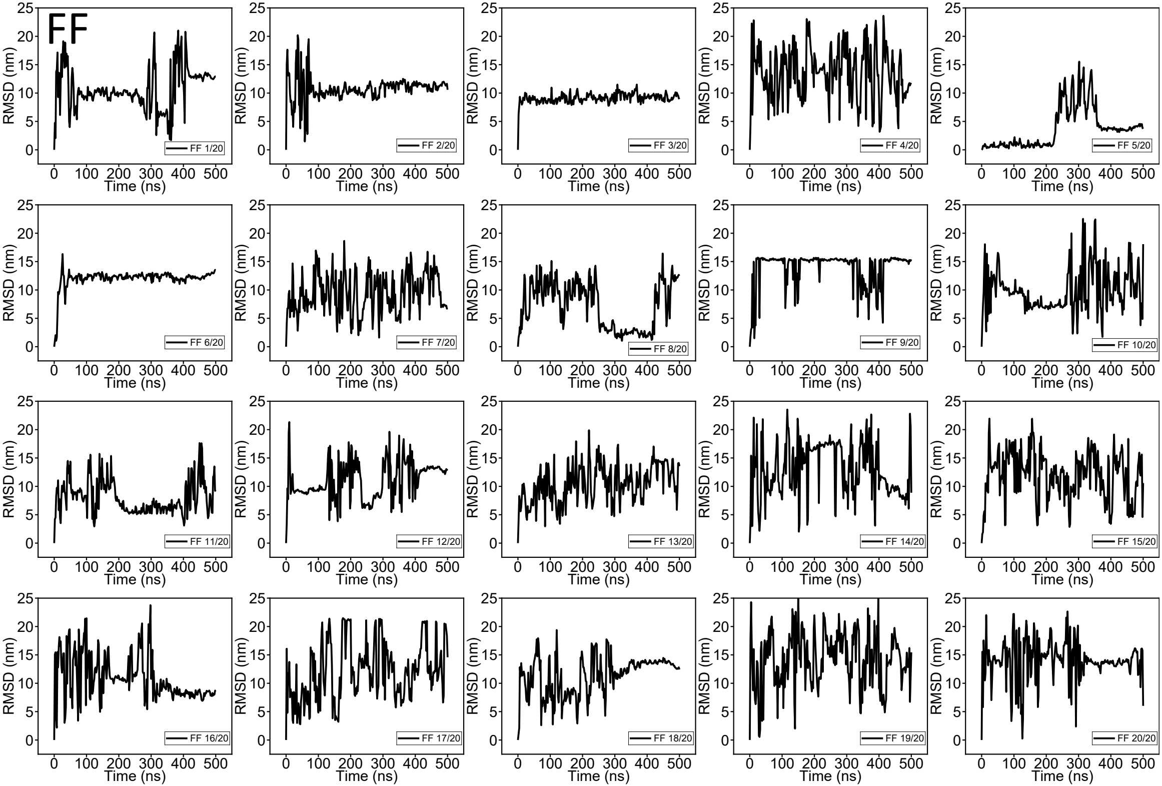

FF
RMSD (nm)
Time (ns)
FF 1/20
FF 2/20
FF 3/20
FF 4/20
FF 5/20
FF 6/20
FF 7/20
FF 8/20
FF 9/20
FF 10/20
FF 11/20
FF 12/20
FF 13/20
FF 14/20
FF 15/20
FF 16/20
FF 17/20
FF 18/20
FF 19/20
FF 20/20

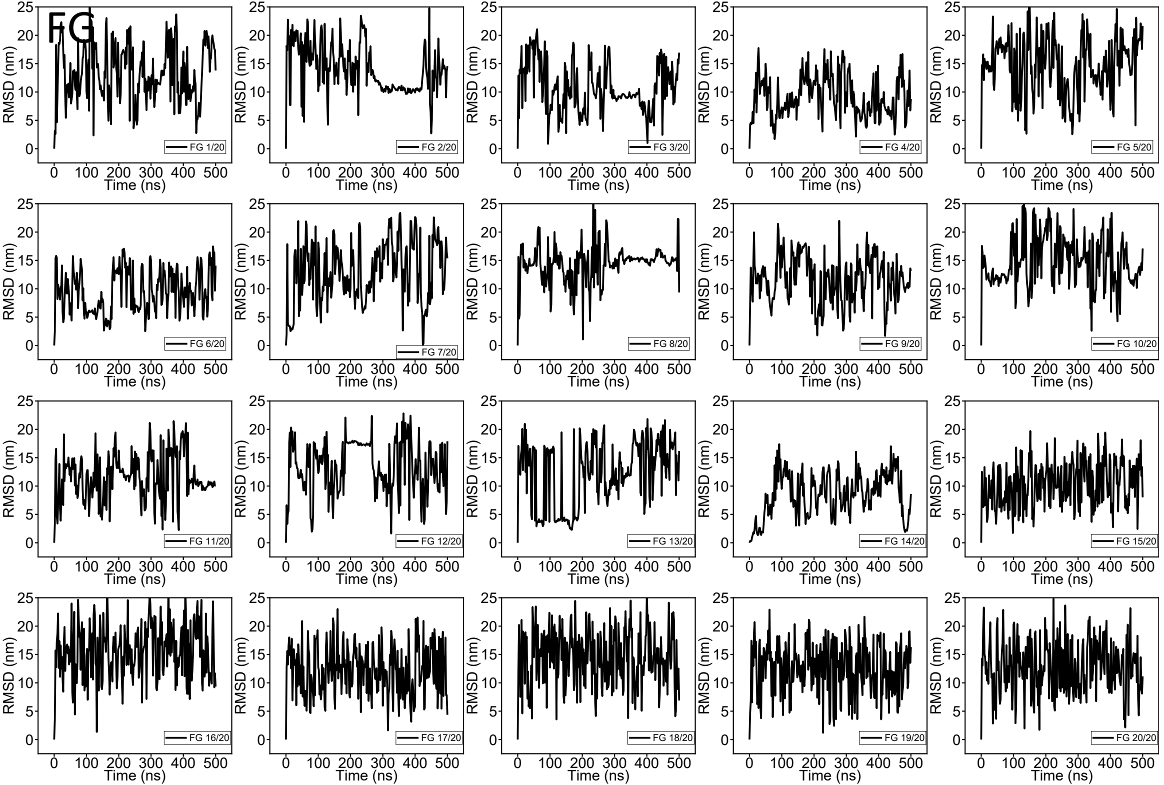
FG
RMSD (nm)
Time (ns)
FG 1/20
FG 2/20
FG 3/20
FG 4/20
FG 5/20
FG 6/20
FG 7/20
FG 8/20
FG 9/20
FG 10/20
FG 11/20
FG 12/20
FG 13/20
FG 14/20
FG 15/20
FG 16/20
FG 17/20
FG 18/20
FG 19/20
FG 20/20

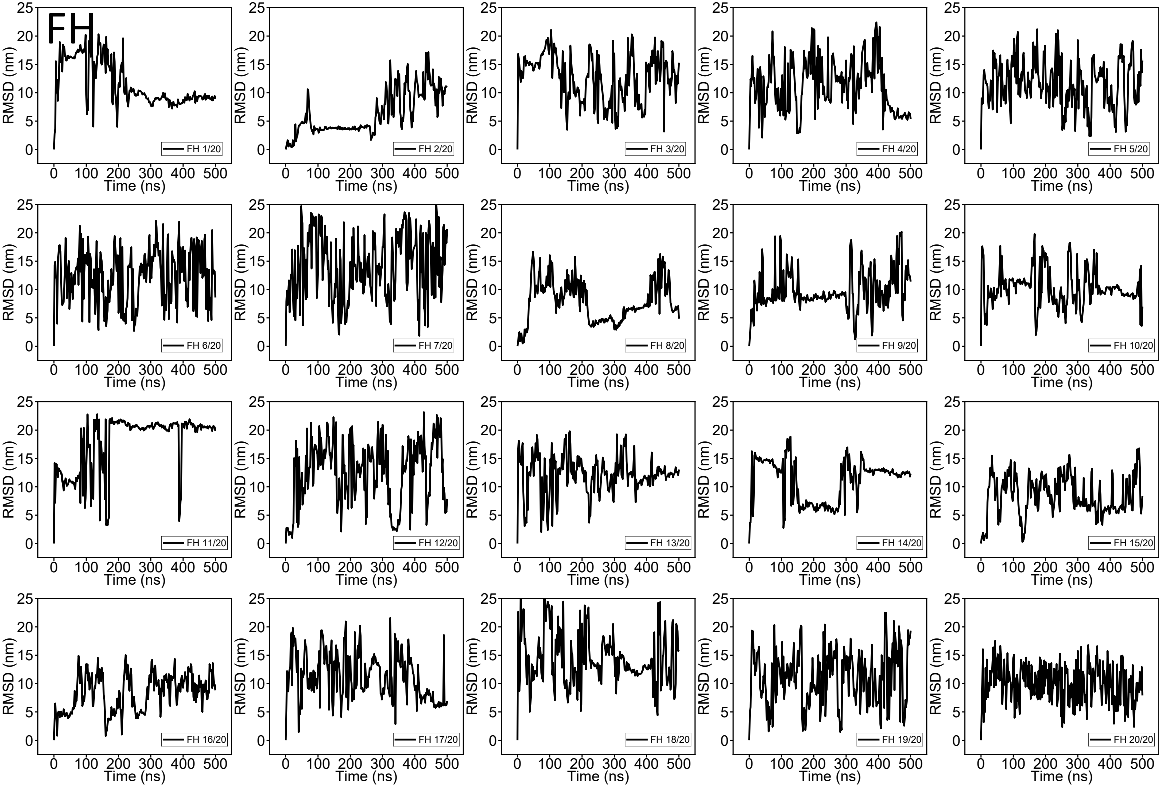
FH
RMSD (nm)
Time (ns)
FH 1/20
FH 2/20
FH 3/20
FH 4/20
FH 5/20
FH 6/20
FH 7/20
FH 8/20
FH 9/20
FH 10/20
FH 11/20
FH 12/20
FH 13/20
FH 14/20
FH 15/20
FH 16/20
FH 17/20
FH 18/20
FH 19/20
FH 20/20

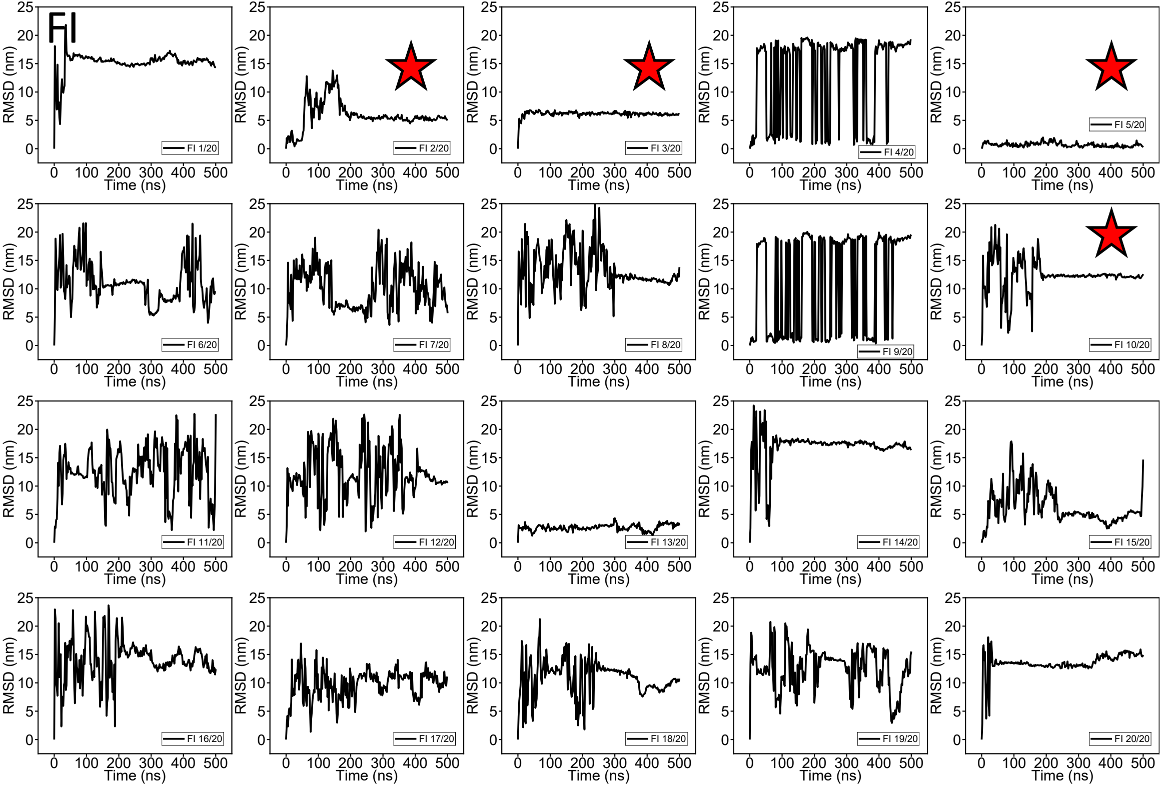
FI
RMSD (nm)
Time (ns)
FI 1/20
FI 2/20
FI 3/20
FI 4/20
FI 5/20
FI 6/20
FI 7/20
FI 8/20
FI 9/20
FI 10/20
FI 11/20
FI 12/20
FI 13/20
FI 14/20
FI 15/20
FI 16/20
FI 17/20
FI 18/20
FI 19/20
FI 20/20

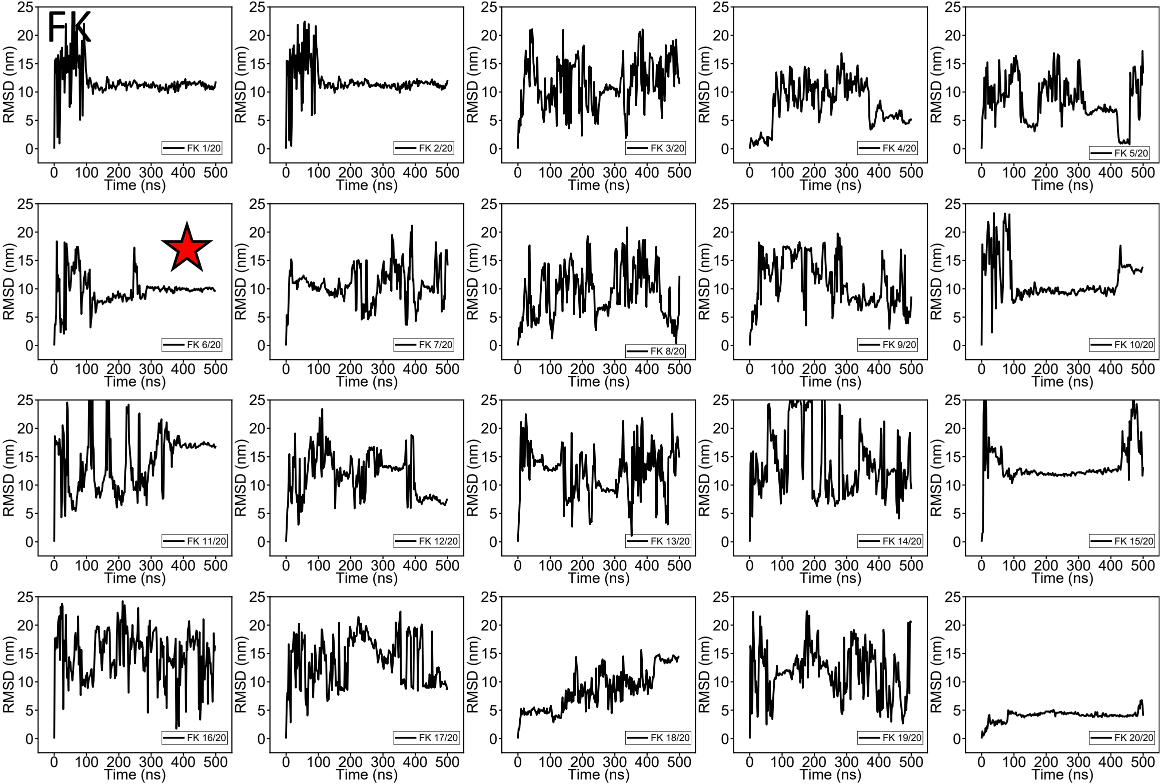
FK
RMSD (nm)
Time (ns)
FK 1/20
FK 2/20
FK 3/20
FK 4/20
FK 5/20
FK 6/20
FK 7/20
FK 8/20
FK 9/20
FK 10/20
FK 11/20
FK 12/20
FK 13/20
FK 14/20
FK 15/20
FK 16/20
FK 17/20
FK 18/20
FK 19/20
FK 20/20

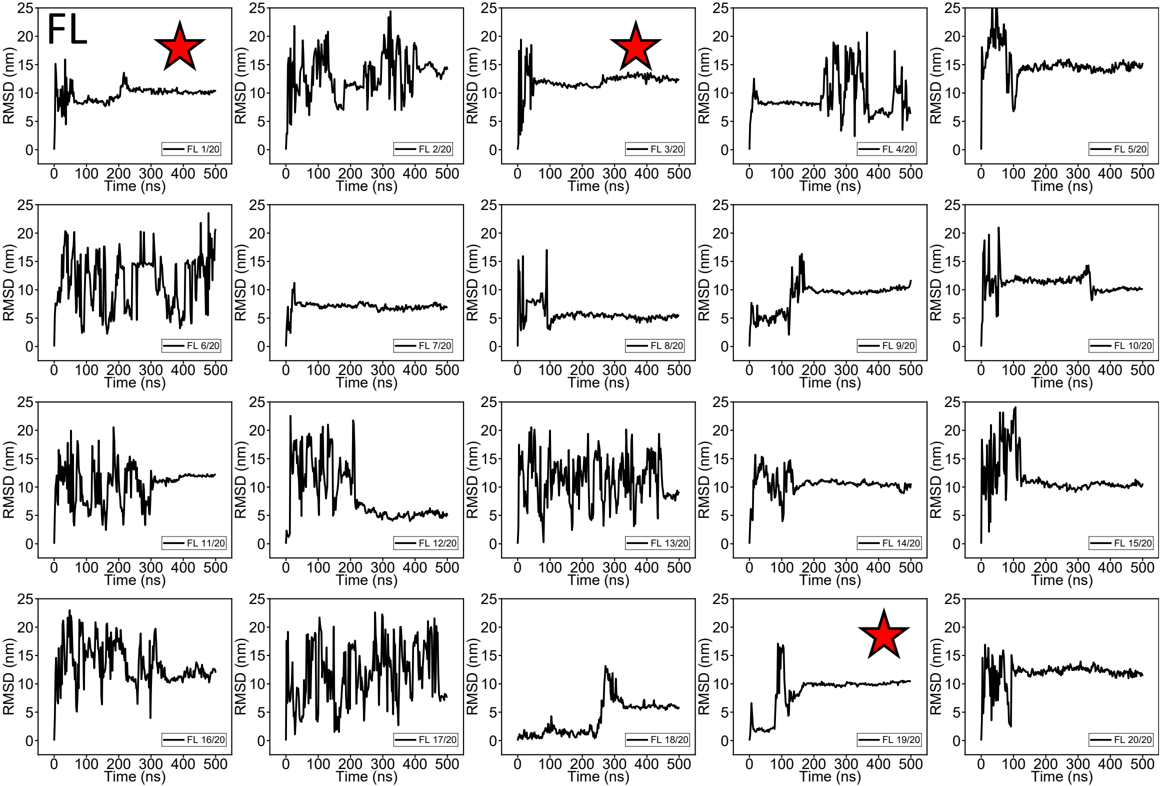
FL
RMSD (nm)
Time (ns)
FL 1/20
FL 2/20
FL 3/20
FL 4/20
FL 5/20
FL 6/20
FL 7/20
FL 8/20
FL 9/20
FL 10/20
FL 11/20
FL 12/20
FL 13/20
FL 14/20
FL 15/20
FL 16/20
FL 17/20
FL 18/20
FL 19/20
FL 20/20

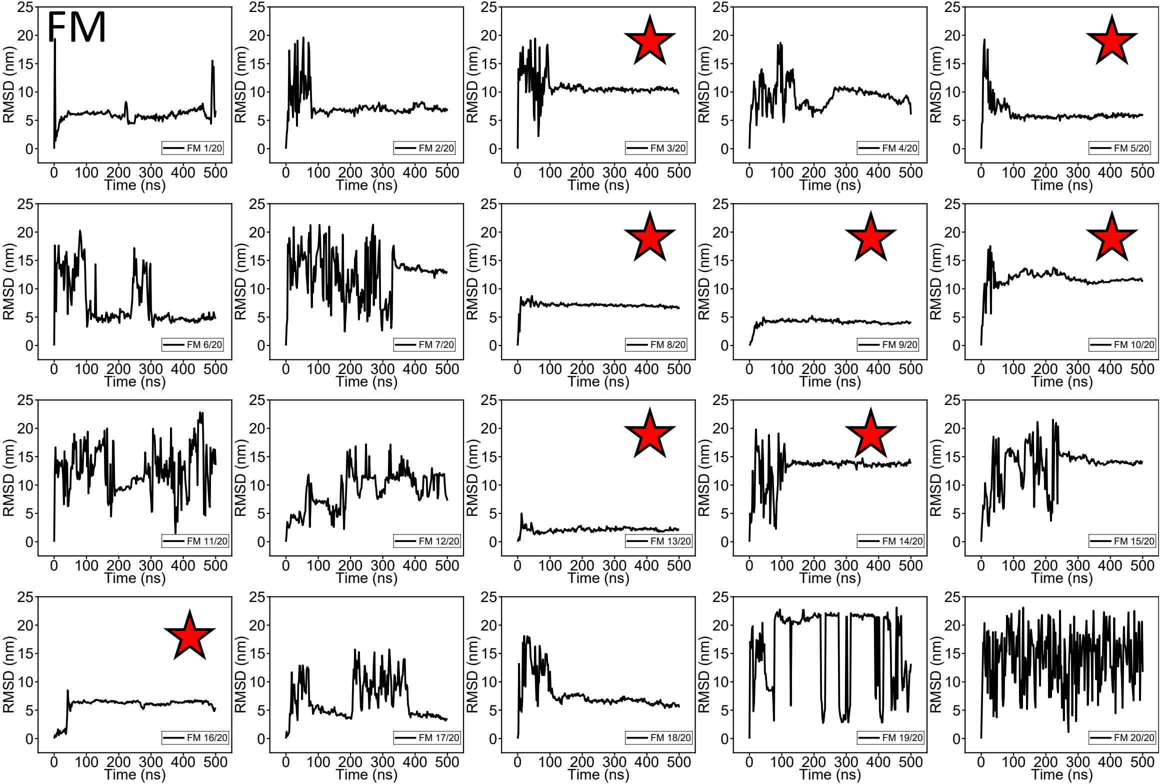
FM
RMSD (nm)
Time (ns)
FM 1/20
FM 2/20
FM 3/20
FM 4/20
FM 5/20
FM 6/20
FM 7/20
FM 8/20
FM 9/20
FM 10/20
FM 11/20
FM 12/20
FM 13/20
FM 14/20
FM 15/20
FM 16/20
FM 17/20
FM 18/20
FM 19/20
FM 20/20

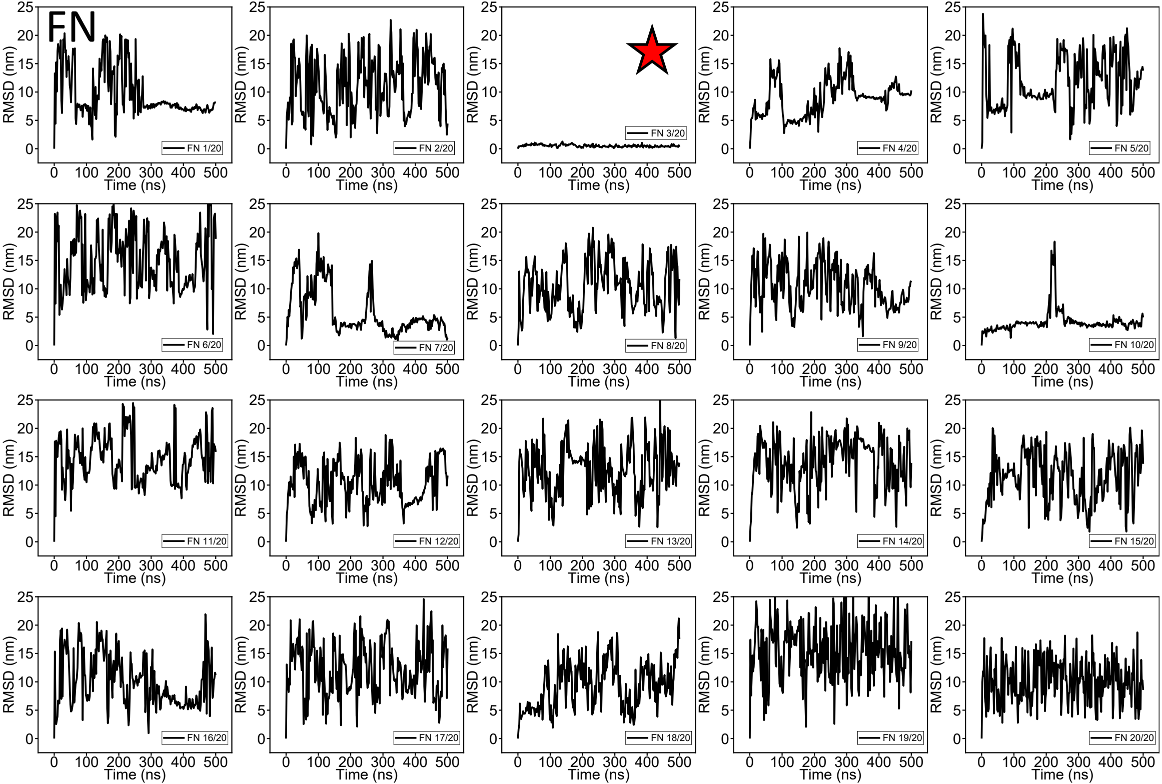

FN
RMSD (nm)
Time (ns)
FN 1/20
FN 2/20
FN 3/20
FN 4/20
FN 5/20
FN 6/20
FN 7/20
FN 8/20
FN 9/20
FN 10/20
FN 11/20
FN 12/20
FN 13/20
FN 14/20
FN 15/20
FN 16/20
FN 17/20
FN 18/20
FN 19/20
FN 20/20

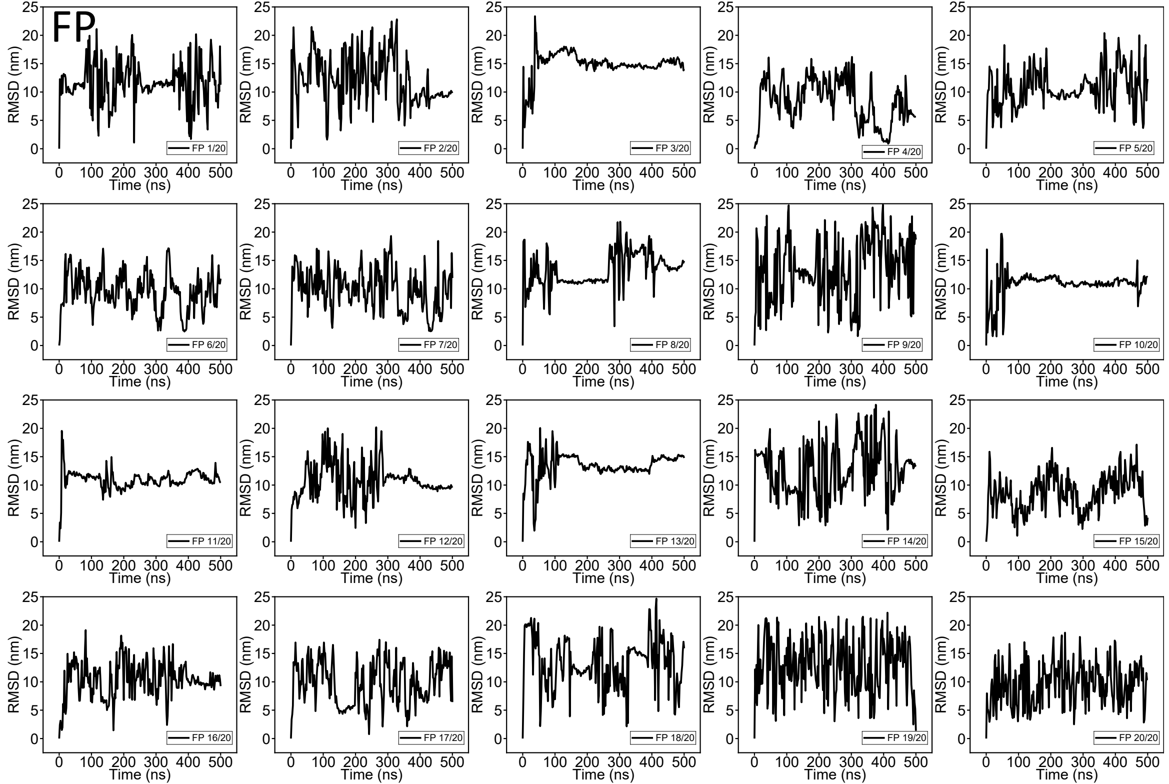
FP
RMSD (nm)
Time (ns)
FP 1/20
FP 2/20
FP 3/20
FP 4/20
FP 5/20
FP 6/20
FP 7/20
FP 8/20
FP 9/20
FP 10/20
FP 11/20
FP 12/20
FP 13/20
FP 14/20
FP 15/20
FP 16/20
FP 17/20
FP 18/20
FP 19/20
FP 20/20

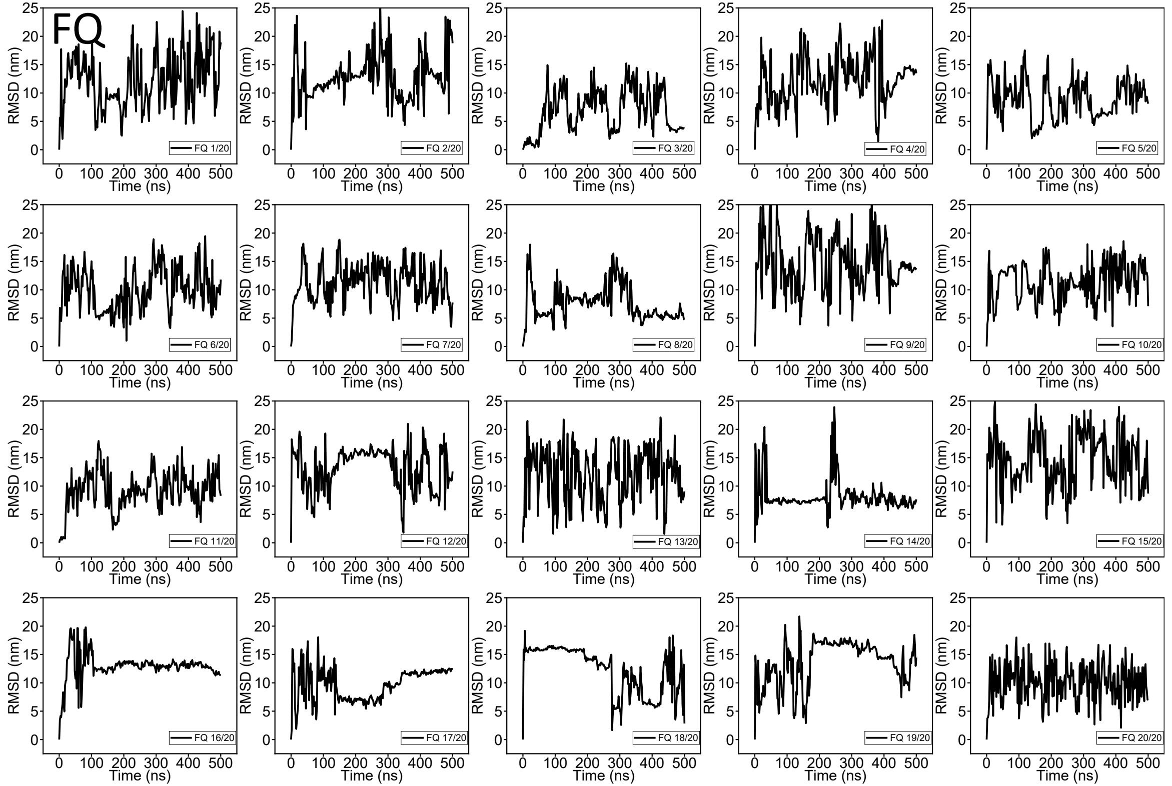
FQ
RMSD (nm)
Time (ns)
FQ 1/20
FQ 2/20
FQ 3/20
FQ 4/20
FQ 5/20
FQ 6/20
FQ 7/20
FQ 8/20
FQ 9/20
FQ 10/20
FQ 11/20
FQ 12/20
FQ 13/20
FQ 14/20
FQ 15/20
FQ 16/20
FQ 17/20
FQ 18/20
FQ 19/20
FQ 20/20

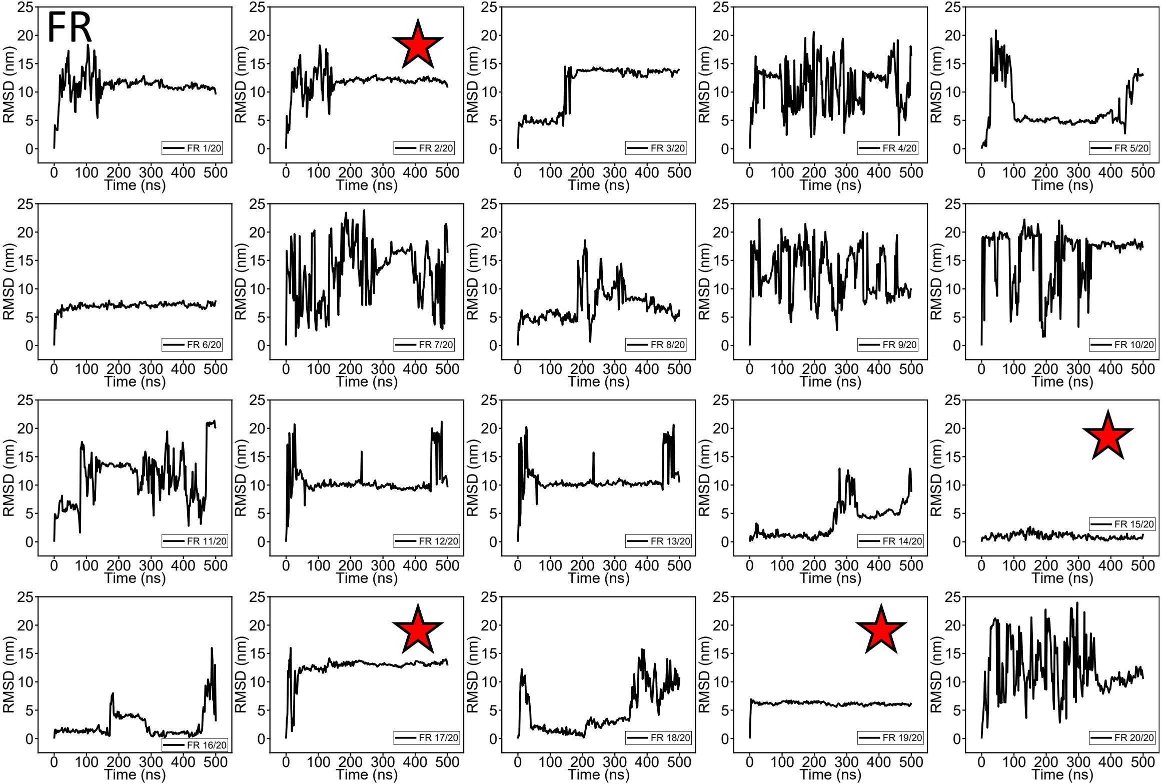
FR
RMSD (nm)
Time (ns)
FR 1/20
FR 2/20
FR 3/20
FR 4/20
FR 5/20
FR 6/20
FR 7/20
FR 8/20
FR 9/20
FR 10/20
FR 11/20
FR 12/20
FR 13/20
FR 14/20
FR 15/20
FR 16/20
FR 17/20
FR 18/20
FR 19/20
FR 20/20

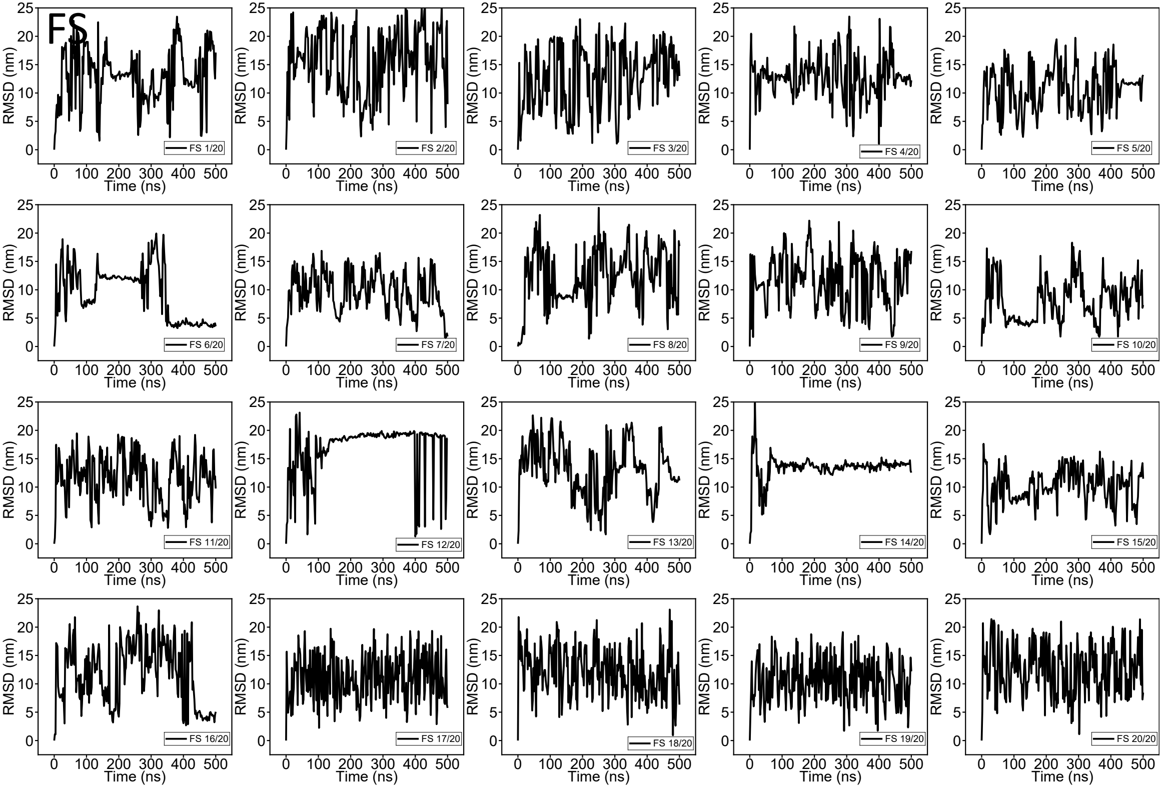
FS
RMSD (nm)
Time (ns)
FS 1/20
FS 2/20
FS 3/20
FS 4/20
FS 5/20
FS 6/20
FS 7/20
FS 8/20
FS 9/20
FS 10/20
FS 11/20
FS 12/20
FS 13/20
FS 14/20
FS 15/20
FS 16/20
FS 17/20
FS 18/20
FS 19/20
FS 20/20

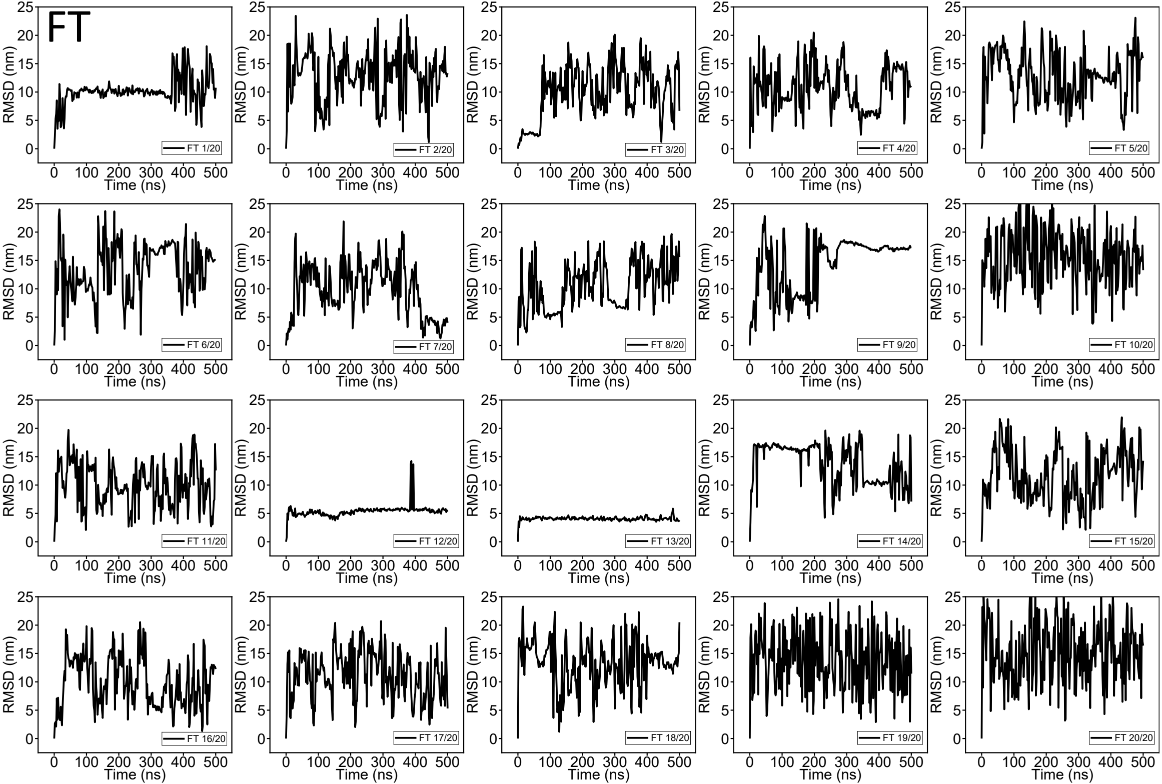
FT
RMSD (nm)
Time (ns)
FT 1/20
FT 2/20
FT 3/20
FT 4/20
FT 5/20
FT 6/20
FT 7/20
FT 8/20
FT 9/20
FT 10/20
FT 11/20
FT 12/20
FT 13/20
FT 14/20
FT 15/20
FT 16/20
FT 17/20
FT 18/20
FT 19/20
FT 20/20

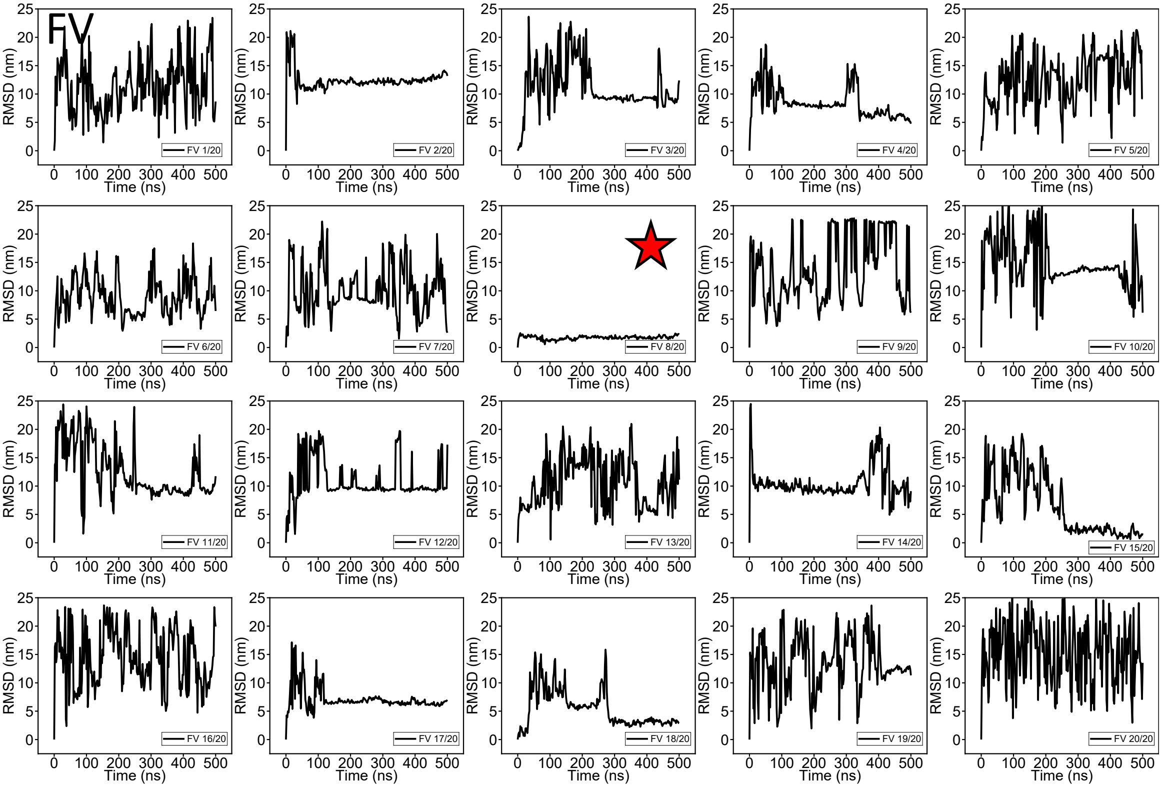
FV
RMSD (nm)
Time (ns)
FV 1/20
FV 2/20
FV 3/20
FV 4/20
FV 5/20
FV 6/20
FV 7/20
FV 8/20
FV 9/20
FV 10/20
FV 11/20
FV 12/20
FV 13/20
FV 14/20
FV 15/20
FV 16/20
FV 17/20
FV 18/20
FV 19/20
FV 20/20

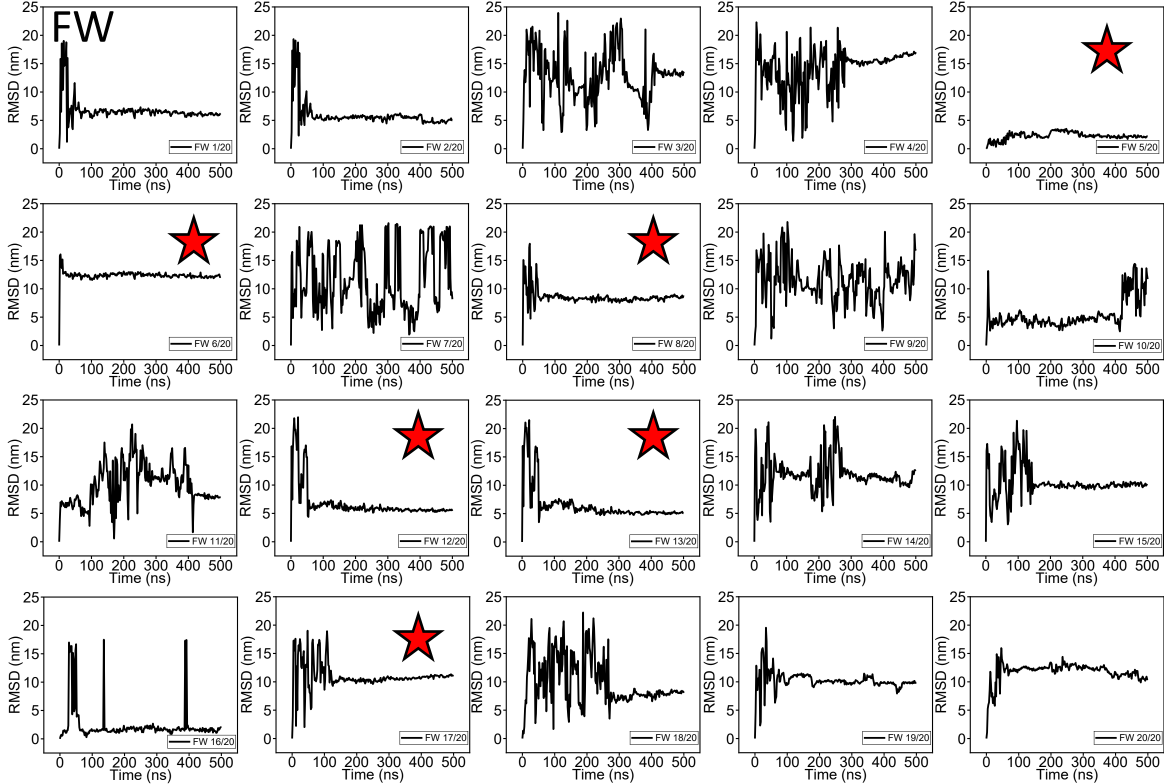
FW
RMSD (nm)
Time (ns)
FW 1/20
FW 2/20
FW 3/20
FW 4/20
FW 5/20
FW 6/20
FW 7/20
FW 8/20
FW 9/20
FW 10/20
FW 11/20
FW 12/20
FW 13/20
FW 14/20
FW 15/20
FW 16/20
FW 17/20
FW 18/20
FW 19/20
FW 20/20

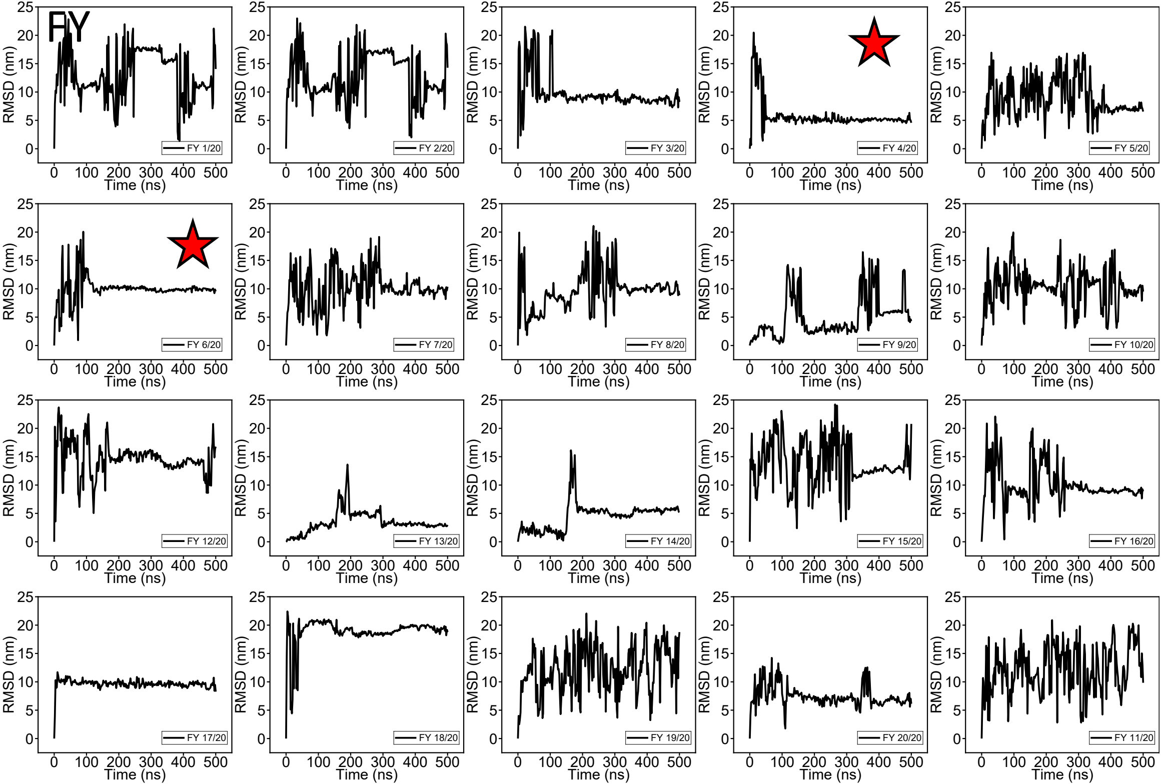
FY
RMSD (nm)
Time (ns)
FY 1/20
FY 2/20
FY 3/20
FY 4/20
FY 5/20
FY 6/20
FY 7/20
FY 8/20
FY 9/20
FY 10/20
FY 12/20
FY 13/20
FY 14/20
FY 15/20
FY 16/20
FY 17/20
FY 18/20
FY 19/20
FY 20/20
FY 11/20
